# Dopamine neurons that inform *Drosophila* olfactory memory have distinct, acute functions driving attraction and aversion

**DOI:** 10.1101/2022.11.23.517775

**Authors:** Farhan Mohammad, Yishan Mai, Joses Ho, Xianyuan Zhang, Stanislav Ott, James Charles Stewart, Adam Claridge-Chang

**Affiliations:** Program in Neuroscience and Behavioural Disorders, Duke-NUS Medical School, Singapore; Institute for Molecular and Cell Biology, A*STAR, Singapore; Department of Pharmacology, National University of Singapore, Singapore; Department of Physiology, National University of Singapore, Singapore; Division of Biological and Biomedical Sciences, College of Health & Life Sciences, Hamad Bin Khalifa University, Qatar

**Keywords:** Motivation, Reward, Valence, Reinforcement, Locomotion, Synapse, Dopamine

## Abstract

The brain must guide immediate responses to beneficial and harmful stimuli while simultaneously writing memories for future reference. While both immediate actions and reinforcement learning are instructed by dopamine, how dopaminergic systems maintain coherence between these two reward functions is unknown. Through optogenetic activation experiments, we showed that the dopamine neurons that inform olfactory memory in *Drosophila* have a distinct, parallel function driving attraction and aversion (valence). Sensory neurons required for olfactory memory were dispensable to dopaminergic valence. A broadly projecting set of dopaminergic cells had valence that was dependent on dopamine, glutamate, and octopamine. Similarly, a more restricted dopaminergic cluster with attractive valence was reliant on dopamine and glutamate; flies avoided opto-inhibition of this narrow subset, indicating the role of this cluster in controlling ongoing behavior. Dopamine valence was distinct from output-neuron opto-valence in locomotor pattern, strength, and polarity. Overall our data suggest that dopamine’s acute effect on valence provides a mechanism by which a dopaminergic system can coherently write memories to influence future responses while guiding immediate attraction and aversion.

## Introduction

For an animal to survive and thrive, its brain must integrate sensory stimuli and internal signals to guide it toward benefits and away from harm. Some neural information has evolved to be innately instructive to behavior—for example, a sensory response to painful heat. Other information has no inherent evolutionary imperative *a priori*, but can acquire behavioral meaning through learning—for example, a naively neutral odor. A fundamental aspect of all brain states is their propensity to make an animal approach or avoid a stimulus, a property termed ‘emotional valence’ [1]. In humans, an emotional behavior like a facial expression of disgust is characterized as having negative valence, while a happy smile can be said to have positive valence. It has long been appreciated that such emotional behaviors have counterparts in all animals, including insects [1–3].

In the brain, emotional valence is partly governed by neuromodulators, which are soluble factors that modify neuronal excitability and synaptic dynamics through their action on metabotropic receptors [4]. Through these cellular effects, neuromodulators transform circuit dynamics, eliciting various motor outputs from a single network [5].

One particularly important neuromodulator is dopamine. In mammals, dopamine-releasing cells are implicated in diverse processes that include motor function, motivation, associative learning and acute valence [6–8]. Many of these functions are conserved across animal species, including the experimentally-tractable vinegar fly, *Drosophila melanogaster*.

In *Drosophila,* dopamine’s importance has mainly been examined in associative functions including olfactory conditioning [9–11], aversive learning [12,13], and memories formed within the mushroom body (MB) [9,10,13–17]. In addition to its diverse roles in olfactory memory [9,11,12,14,18–28], dopamine in the MB has also been implicated in regulating various non-olfactory, non-memory behaviors in *Drosophila.* These include innate odor preference [29–31], visual learning [32], sleep and circadian rhythm [33,34], temperature preference [35], oviposition choice [36], courtship drive [37], wing-extension during courtship [38], decision-making [39,40], visual-attention [41], maintenance of flight state [42], odor discrimination [43], sensitivity to odor cues [44,45], discrimination and reaction to novel space [46], as well as signaling satiety and hunger to control foraging behavior [47–49]. Other recent studies have shown that dopamine modulates sensory processing. For example, dopamine release in the antennal lobe (the primary olfactory processing center in *Drosophila*) enhances odor discrimination [43], and also regulates the activity of local inhibitory neurons, leading to increased activity in projection neurons and enhanced sensitivity to odor cues [44,45].

The synaptic fields of the MB are formed from the confluence of ∼2000 odor-responsive sensory Kenyon cells (KCs), 34 mushroom-body output neurons (MBONs), and ∼120 dopaminergic cells (DANs) [50]. Both DANs and KCs synapse with MBONs [51], which affect behavioral valence by influencing locomotion [52]. Changes in olfactory valence arise when DANs modulate KC→MBON synaptic strength via dopamine release [53–55], resulting in the assignment of negative or positive valence to an odor response when coincident events activate aversive or appetitive DANs, respectively [55].

Notably, DANs have also recently been associated with an array of different valent stimuli and behavior. DANs in the paired-anterior-medial (PAM) cluster, which drives appetitive associative olfactory learning [9,26], innately respond to positively valent odors and are associated with upwind orientation and approach in response to those odors [45,56–59]. They are also implicated in responses to sugars and food-seeking behaviors [44,57,58,60,61], place-preference tasks [62,63], and response to carbon dioxide in larvae [64]. These studies have, however, largely examined the role of PAM DANs with respect to external stimuli, with less examination of their intrinsic, ongoing effects on behavior. When activated with thermogenetic or optogenetic channels, MBONs can autonomously drive valent behavior—even in the absence of odorants, food, or other orienting sensory stimuli [52]. This phenomenon of MBON valence establishes that MB circuits are capable of driving non-associative, acute valence. As such, an important question goes unanswered: is DAN activity in isolation also capable of driving acute valence behavior? To our knowledge, only two other studies have explored this question explicitly, demonstrating heterogeneous DAN-mediated acute valence [51,65].

In the present study, we addressed this question using optogenetic experiments, in which freely moving flies bearing optogenetic constructs were allowed to approach or avoid artificial activation of genetically defined dopaminergic cells in the PAM cluster. We found that flies can be attracted to or repelled by activation in some DANs, but are largely indifferent to activation in others. Genetic lesions indicate that dopaminergic acute valence is independent of MB sensory and associative functions, suggesting that this behavior is distinct from learning and that PAM activity is capable of driving valence in the absence of external stimuli. In a broad driver, valence depends on dopamine, glutamate and octopamine; in the β-lobe DANs, valence depends on both dopamine and glutamate, establishing roles for co-transmitters. An optogenetic inhibition experiment of the β-lobe DANs revealed that, even in a low-stimulus environment, pre-existing neural activity contributes to ongoing locomotor behavior.

## Results

### Mushroom-body dopamine neurons drive approach and avoidance

To investigate the role of dopamine on acute approach/avoidance behaviors, we generated transgenic flies expressing an optogenetic activator (Chrimson, henceforth ‘Chr’) [66] in the PAM dopaminergic neurons (PAM DANs) that project to the MB [9,10,67]. The flies were then analyzed in a light-dark choice assay (**Figure 1a**). We selected *R58E02*—a transgenic line of a flanking region of the *Dopamine transporter* (*DAT*) gene fused with *Gal4* [68,69]. This driver is expressed in a large subset of the PAM DANs, and has fibers in a broad set of MB neuropil zones [9,10,24,25,60,70] (**Figure 1b**, **S1a, Movie S1**). Valence was calculated as the mean difference between optogenetic test flies and the corresponding controls, and displayed as effect-size curves [71]. Flies expressing the Chr opto-activator with the *R58E02* transgene were strongly attracted to light (**Figure 1e**, **Movie S2**). By contrast, flies expressing another driver, *R15A04*, which is expressed in PAM DANs that send fibers to a more restricted set of MB zones [25,60,69] (**Figure 1c**, **S1b, Movie S3**) tended to avoid the light at the highest illumination intensity (70 µM/mm^2^, **Figure 1f**).

**Figure 1.**
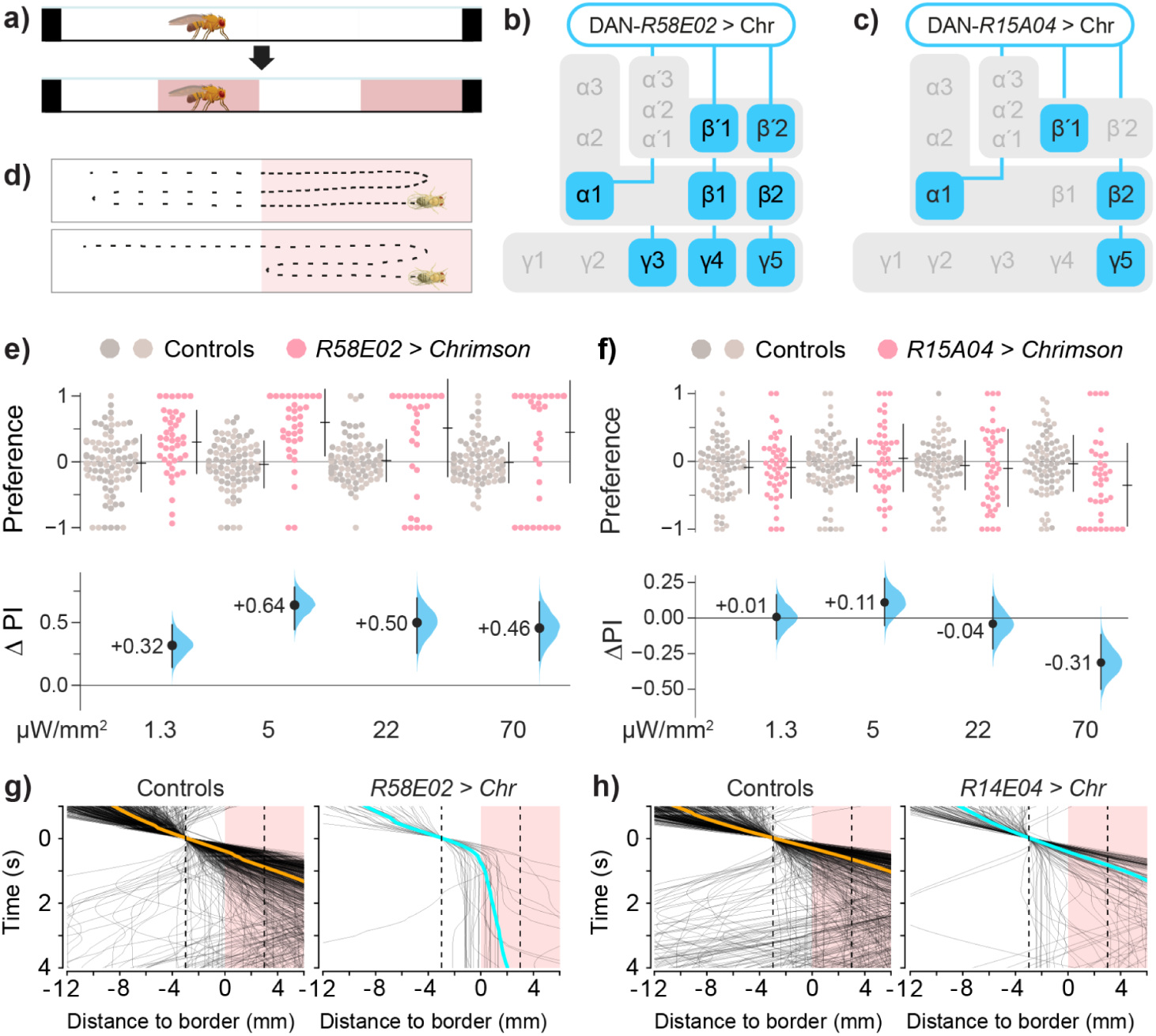
Activities in different dopaminergic cells drive valence. **a.** Schematic of the optogenetic assay showing that after an initial dark phase, half of the chamber is illuminated with two bands of red light. See Methods for further details. **b–c.** Schematic of the two DAN drivers, R58E02 and R15A04, with projections to MB synaptic zones. R58E02 is expressed in nearly all PAM types, projecting to α1, β1, β2, β’1, β’2, γ4, and γ5, with weaker expression in γ1, γ2, and the peduncle. R15A04 is expressed in PAMs that project to the α1, β2, β’1, and γ5 zones. **d.** Schematic of the hypothetical locomotor modes for valence. **Top** Flies move slowly in the favored area. **Bottom** Flies maneuver to remain in the favored area. Either mode increases the time spent in the preferred area. **e.** R58E02>Chr flies spent more time in the light zones. The **upper panel** shows the preference indices (PIs) for test flies (red dots) and driver and responder controls (R58E02/+ and Chr/+, gray dots). The **lower panel** shows the valence effect sizes (mean differences, ΔPI) between control and test flies, with confidence intervals (black line) and the distribution of ΔPI error (blue curve). The positive ΔPI values indicate a positive valence. See **Table S1** for detailed genotypes and statistics. **f.** *R15A04>Chr* flies avoided opto-activation. The negative ΔPI values indicate avoidance. **g.** Walking behavior of the subset of flies that entered the choice zone from the dark side and approached the dark-light interface. Only data from flies that approached the choice zone were included. Traces of *R58E02>Chr* paths (black) are aligned to choice-zone entry, i.e. locked to the time of entering the boundary area. The colored lines show the overall mean trajectory. The horizontal axis is aligned to the middle of the choice zone. Test flies slowed or stopped at the boundary,
with their heads on either side of the middle of the light interface. **h.** Traces of *R15A04>Chr* flies as they entered the choice zone from the dark side. Trajectory data were taken from epochs with 70 μW/mm illumination. Data for all panels can be found in the corresponding folder on the Zenodo data repository (https://doi.org/10.5281/zenodo.7747425).

Thus, while activation of the *R58E02* DANs drives strong positive valence, flies will avoid activation in the subset defined by *R15A04*.

### DAN activation influences locomotion

When traversing a boundary between two stimulus areas, walking flies encountering aversive stimuli use reversals or turns to maneuver away [14,52]. A fly that displays a spatial preference for one of two areas, however, could hypothetically employ another locomotor mode: slowing down in the favored area (**Figure 1d**). We thus explored how valence, choice, and speed were associated in *R58E02* and *R15A04* flies. Regressions between preference, speed ratio, and a choice index showed that in these lines, preference was more strongly determined by differential speed (**Figure S1c-f**). We also inspected locomotion at the boundary by aligning the dark-to-light trajectories in a single experiment. In the subset of flies that entered the boundary choice-zone, trajectory lines drawn over time indicated that as the optogenetic DAN lines traverse into the choice zone from the dark side, the *R58E02>Chr* flies tend to walk slower and frequently stop in the boundary area (**Figure 1g-h**). These observations indicate a relationship between DAN valence and changes in walking speed.

### Olfactory circuits are dispensable to broad DAN attraction

We next investigated whether circuit and molecular components that are essential for memory formation are similarly necessary for valence. We started by investigating the KCs, as DANs instruct odor memory by modulating KC function [53,55]. The expression of a conditioned response in olfactory associative learning (that is, avoidance of or approach towards a conditioned odor) is also reliant on KC activity [72,73]. We asked whether odor memories could be impaired by expressing the light-actuated anion channelrhodopsin, GtACR1 (hereon ‘ACR1’) in the KCs using the *MB247* driver and actuating with green light. As hypothesized, in aversive shock–odor conditioning with ACR1 actuation (**Figure 2a**), flies expressing the opto-inhibitor in the KCs (*MB247>ACR1*) failed to learn (**Figure 2b**). This finding demonstrates that ACR1 sufficiently inhibits KCs to abolish memory formation. In *R58E02>Chr* flies, green light successfully induced synthetic, optogenetic appetitive memory; however, in *R58E02>Chr, MB247>ACR1* flies, light actuation did not induce synthetic memory (**Figure 2c**), further verifying that ACR1 elicits KC inhibition and can block the memory-inducing effect of *R58E02* dopamine cells. We thus demonstrated the successful inhibition of KCs using the opto-inhibitor ACR1, with functional consequences for memory acquisition both using a real stimulus, electric shock, and optogenetic stimulation of the PAMs.

**Figure 2:**
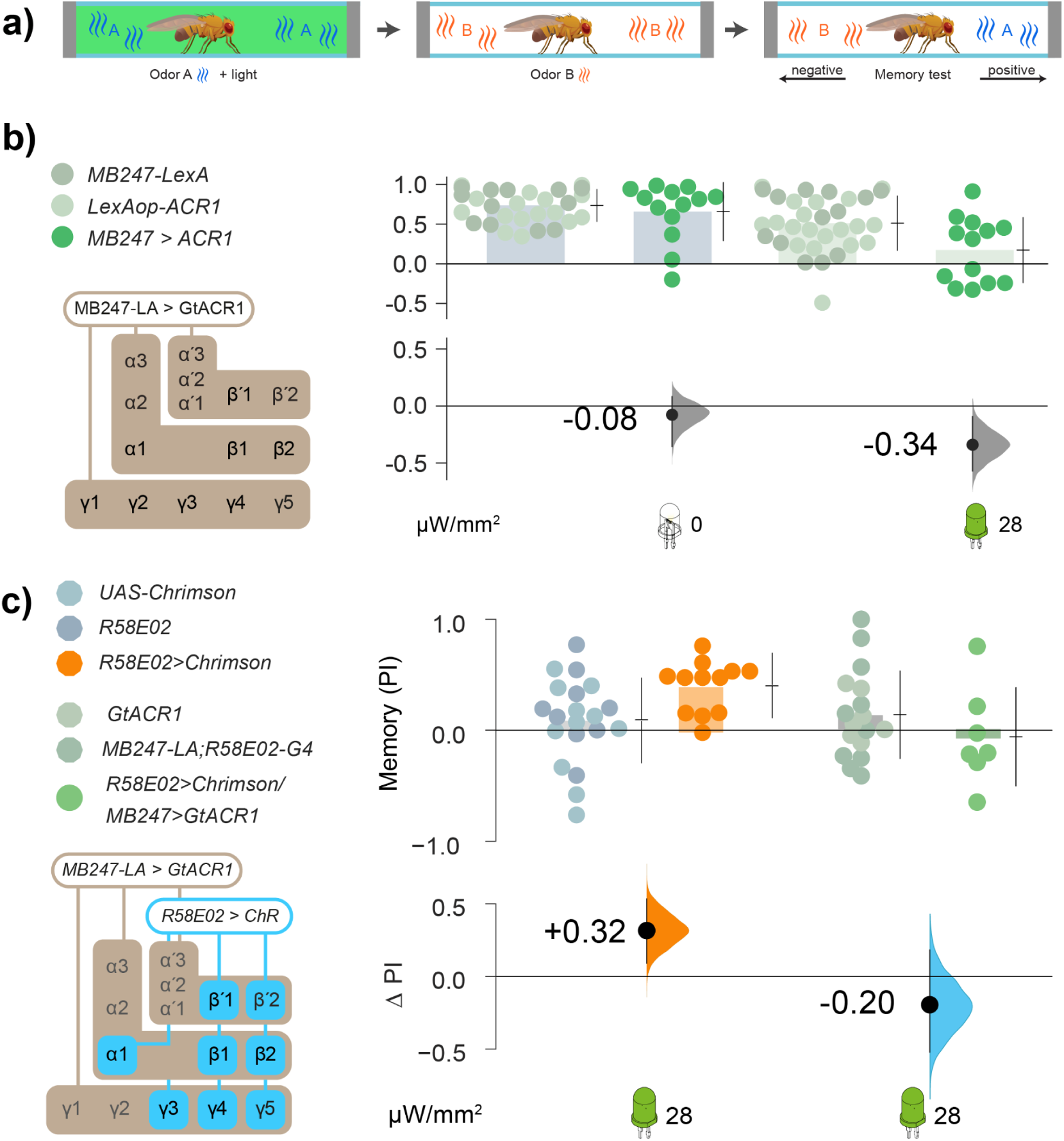
Inhibition of KC activity with GtACR1 prevents the formation of shock-conditioned and DAN-conditioned olfactory memories. **a.** Optogenetic olfactory conditioning protocol: two odors were presented in sequence, one paired with green light; the two odors were then presented in different arms of the chamber to test the conditioned preference (see Methods for details). **b.** Silencing the KCs with ACR1 actuation decreased aversive shock memory by 75.0% [95CI-44%, −104%]. The genetic controls were *MB247-LexA/+* and *LexAop-ACR1/+* (gray dots); the test animals were *MB247-LexA/LexAop-ACR1* (green dots). Unilluminated flies (both control and test animals) showed robust learning with shock-paired odor (**left side**, 0 µW/mm^2^). Illumination with green light reduced the PI of test flies by ΔPI = –0.34 relative to the genetic controls (**right side**, 28 µW/mm^2^). **c.** Activation of DANs with green light paired with odor, instructed an attractive olfactory memory in *R58E02>Chr* test flies (orange dots) relative to controls (*R58E02-Gal4*/+ and *UAS-Chr/*+, **left side** gray dots). The contrast between test and control animals is ΔPI = +0.32 (orange curve). Green light induced almost no memory formation in *R58E02>Chr; MB247>ACR1* flies (**right side**, gray dots). The difference attributable to ACR1 inhibition corresponds to a performance reduction of –120% [95CI –200%, –24%], i.e. a shift from attraction to mild aversion. Sample sizes: *N*_*experiments*_ = 6, 6, 6; *N*_flies_ = 144 per genotype. Data for all panels can be found in the corresponding folder on the Zenodo data repository (https://doi.org/10.5281/zenodo.7747425).

We then questioned whether KC activity was likewise required for acute valence. We implemented a strategy to allow flies to simultaneously activate DANs with Chr and inhibit KCs using ACR1 [74,75]. As both channelrhodopsins are responsive to green light, we tested the ability of green light to both activate DANs and silence KCs. *R58E02>Chr* flies were attracted to green light (**Figure 3a**), confirming effective Chr actuation. One possible confound would be if flies responded to KC inhibition with a strong attraction that masked DAN attraction; in a valence test, however, *MB247>ACR1* flies exhibited only mild aversion (**Figure 3b**). We then tested flies carrying the *R58E02>Chr, MB247>ACR1* genotype for their optogenetic preference. Even with inhibited KCs, valence remained intact (**Figure 3c**; **Figure S2ab**).

**Figure 3.**
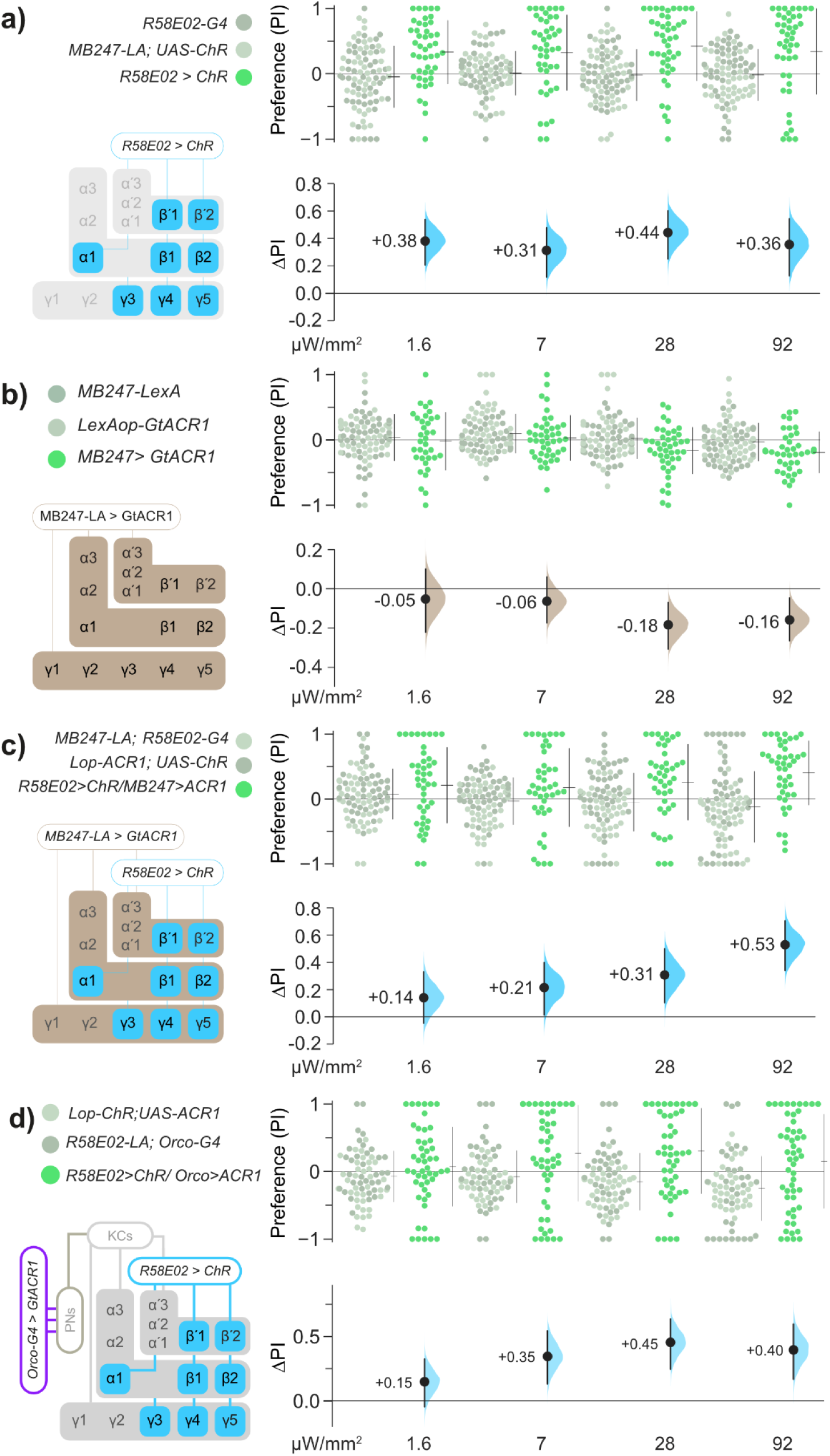
Kenyon cells are dispensable for *R58E02* DAN valence. **a**. *>R58E02>Chr* flies are attracted to green light. The left schematic illustrates the expression pattern of *R58E02*. Flies carrying all three transgenes displayed attraction to green light (green dots), resulting in positive valence (black dots and blue curves in the lower panel). Parental-type control flies (*R58E02-Gal4/+* or *MB247-LexA/+; UAS-Chr/+*, gray dots) showed a neutral preference for green light. **b.** Relative to genetic controls, *MB247-LexA>lexAop-ACR1* flies displayed a modest avoidance of green light at high intensities (22 and 72 µW/mm^2^). The schematic indicates that MB247-LexA drives ACR expression in most MB-intrinsic KCs. **c.** In *R58E02>Chr/MB247>ACR1* flies, preference for DAN activation mediated by *R58E02-Gal4>UAS-Chr* was unaffected by simultaneous opto-inhibition of the MB intrinsic cells with *MB247-LexA>lexAop-ACR1*. Effect sizes (blue curves) show the net effect of comparing test flies carrying all four transgenes (green dots), with controls (gray dots). **d.** Simultaneous inhibition of ORNs using *Orco>ACR1* had little impact on attraction to PAM DAN activation. At 92 μW/mm^2^, the ΔPI was +0.40 [95CI +0.17, +0.60], P = 0.001. Transgene abbreviations: *LexAop = LOP, Gal4 = G4*. Data for all panels can be found in the corresponding folder on the Zenodo data repository (https://doi.org/10.5281/zenodo.7747425).

We also performed experiments that showed that the olfactory receptor neurons (ORNs) are not required for PAM valence. Activating PAMs and simultaneously inhibiting ORNs still resulted in positive valence (**Figure 3d**, **S2c**). These results indicate that for PAM-mediated acute attraction, components of the olfactory circuit required for memory are dispensable for valence at both the primary sensory level (ORNs) and at higher levels of representation (KCs).

### Dopamine receptors in the KCs are inessential for broad PAM attraction

Dopamine-receptor function in the KCs is required for olfactory learning [19,55,76]. Targeted *Dop1R1* knockdown in the KCs almost entirely ablates both short-term and long-term learning, while *Dop1R2* knockdown has an impact only on long-term learning, and *Dop2R* knockdown has no effect on either [77]. To determine the dopamine-receptor KC requirements for *R58E02-*mediated attraction, we knocked down *Dop1R1, Dop1R2,* and *Dop2R* in KCs using RNAi transgenes [78]. In a *R58E02-LexA* > *LexAop-Chr* background, individual knockdowns of *Dop1R1* and *Dop1R2* each caused only minor reductions in *R58E02*-mediated valence, while the knockdown of *Dop2R* resulted in a moderate reduction in valence (–41%) (**Figure 4a–c**). Along with the evidence that KC activity itself is not required, these data showing different receptor dependencies verifies that acute PAM valence and olfactory learning are mediated by distinct mechanisms. Note that the partial dependence of valence on Dop2R contrasts with our finding that acute optogenetic inhibition of KC activity has no effect on *R58E02*-mediated valence, which could be explained by long-term changes as a result of chronic *Dop2R* knockdown.

**Figure 4.**
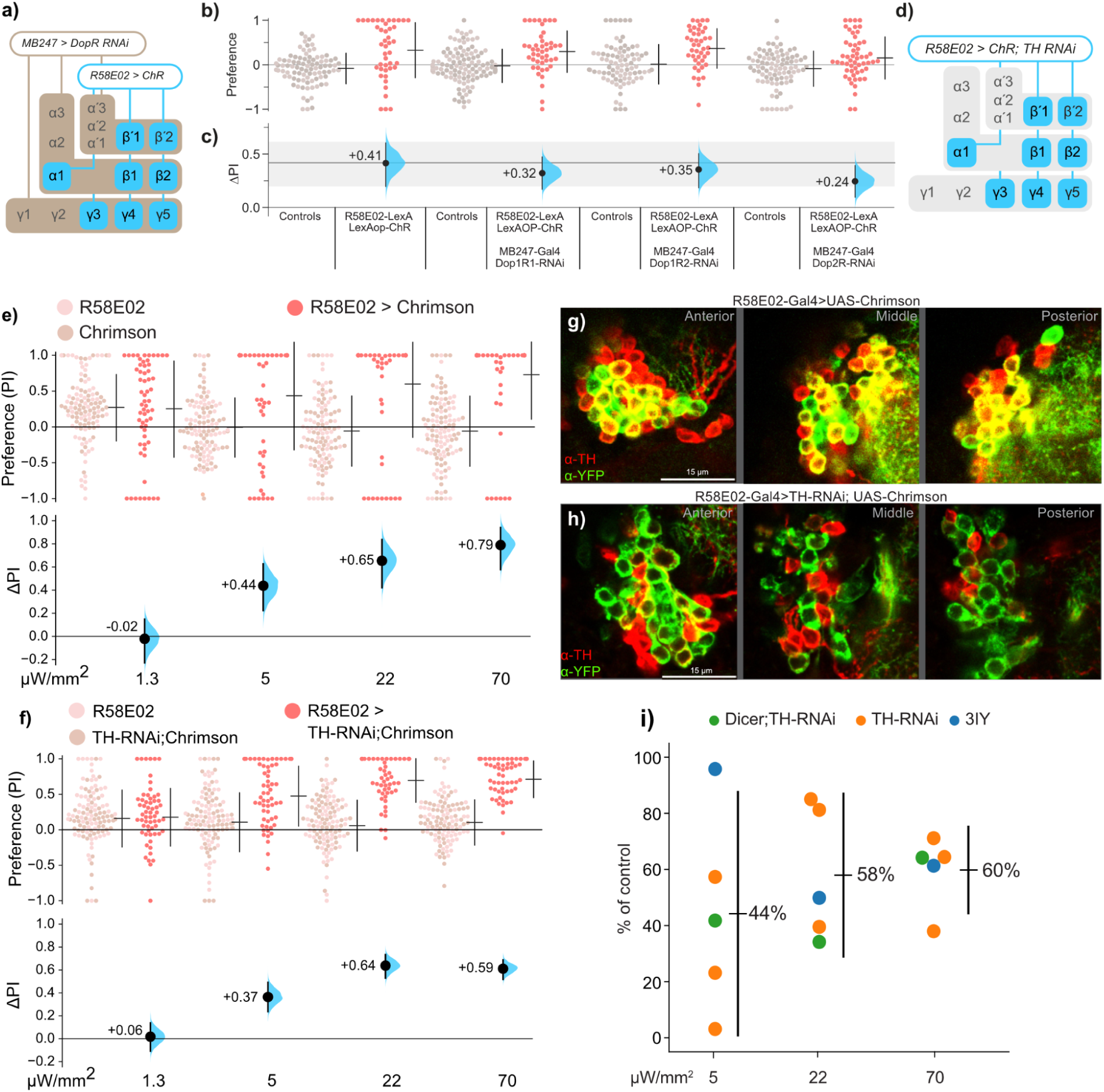
For driving DAN-mediated attraction, dopamine is a partial contributor. **a.** Schematic of the use of *MB247-Gal4* to knockdown receptor expression by RNAi in the *R58E02-LexA>lexAop-Chr* optogenetic background. **b-c.** Knocking down *Dop1R1, Dop1R2*, and *Dop2R* in KCs had minor effects on *R58E02-LexA>lexAop-Chr* light attraction. The gray ribbon indicates the control-valence 95% confidence interval. Experiments used 72 µW/mm^2^ red light. **d.** Schematic for the use of *R58E02-Gal4* to simultaneously express Chr and knockdown TH expression. **e.** Immunohistochemistry of the PAM DAN cluster stained with α-TH (red) and α-YFP (green) in flies expressing Chr-YFP in *R58E02* cells. Yellow rings indicate the co-localization of α-TH and α-YFP signals in cells in the PAM cell-body cluster at three optical slices. **f.** Immunohistochemistry images of the DANs with TH-RNAi co-expression, showing that cells with an α-YFP signal (*R58E02* cells) have a greatly lower α-TH signal. **g-h.** Knocking down TH expression with TH-RNAi has a moderate effect on *R58E02* valence across four intensities. For example, at 70 µW/mm^2^ the valence is +0.79 ΔPI in *R58E02>Chr* flies (G) and is reduced to +0.59 ΔPI in flies carrying the UAS-TH-RNAi knockdown transgene (H). **i.** Averaging summary of the effects of reducing dopamine on *R58E02*-mediated valence with either gene knockdown (*UAS-TH-RNAi*, with or without *UAS-Dicer*) or a chemical inhibitor of TH activity (3-Iodo-L-tyrosine, 3IY). Each dot represents the percentage effect size of light intensity in an experiment (i.e., the *R58E02>Chr; TH-RNAi* experiment was replicated three times). Across all three intensities in five experiments, dopamine depletion resulted in an average ∼46% reduction in valence. The vertical line indicates the 95% confidence interval. Data for all panels can be found in the corresponding folder on the Zenodo data repository (https://doi.org/10.5281/zenodo.7747425).

### Dopamine has a partial role in R58E02 valence

As PAMs are dopaminergic, we expected that depletion of dopamine from the PAMs would result in a loss of most if not all valence. We tested this hypothesis by depleting dopamine in DANs by several methods. First, we used RNAi against tyrosine hydroxylase (TH), an essential enzyme for dopamine synthesis [79] and encoded in flies by the *TH* gene (also referred to as *pale*). Compared to flies with intact *TH* expression (**Figure 4e**), flies with reduced *TH* in the *R58E02* DANs exhibited only a modest valence reduction (**Figure 4f**). This partial valence reduction was observed in the three higher light intensities, and across three replications of this experiment (**Figure 4e-f**, **S3a-d**). To confirm that *TH* knockdown was effective, we performed immunohistochemical staining of the DANs in *R58E02 > TH-RNAi* flies, which showed that TH expression was markedly reduced (**Figure 4g-h**, **Movie S4-S5**).

Second, we aimed to amplify the RNAi transgene’s efficacy with the simultaneous overexpression of Dicer2 endonuclease [78]; this approach resulted in overall valence (e.g. at 70 µW/mm^2^) that was not substantially different from the RNAi alone, i.e. a partial reduction (**Figure S3e, 4i**). Third and finally, we depleted dopamine systemically by feeding flies 3-iodotyrosine (3-IY), a competitive inhibitor of TH [80,81]. This pharmacological intervention also resulted in *R58E02>Chr* valence exhibiting only a partial reduction (**Figure S3f-g**). To gain a summary estimate of the overall effect of removing dopamine from the PAMs on valence, we averaged the results across all three DA-depleting interventions (*TH-RNAi* alone, *TH-RNAi* with *Dicer2*, and 3-IY). Averaging the three interventions in each of the three light intensities indicated that dopamine depletion reduces valence to 44%, 58%, and 60% of control *R58E02*-mediated valence in 5, 22, and 70 mW/mm^2^ illumination, respectively (**Figure 4i**). Averaging across the intensities gives 54% of control levels, i.e., an overall reduction of –46% (**Figure 4i**). Thus, over a range of light conditions and with several different lesions, dopamine appears to mediate roughly half of *R58E02* valence.

### Broad DAN attraction relies on glutamate and octopamine

Single-cell RNA sequencing data have shown that PAMs express neurotransmitter-related genes aside from those pertaining to dopamine [28]. As dopamine did not fully account for *R58E02* valence, we hypothesized that the dopamine-independent component of *R58E02* valence might depend on other neurotransmitters. We knocked down genes involved in the synthesis or vesicular transport of four other transmitters: *vGlut* for glutamate, *Gad1* for GABA, *vAChT* for acetylcholine, and *Tꞵh* for octopamine. Of these, knockdown of *vGlut* and *Tꞵh* in the *R58E02* cells produced effects on valence that were comparable in magnitude to the dopamine reduction: –48% reduction for *Vglut* and –38% for *Tbh* (**Figure 5a**).

**Figure 5.**
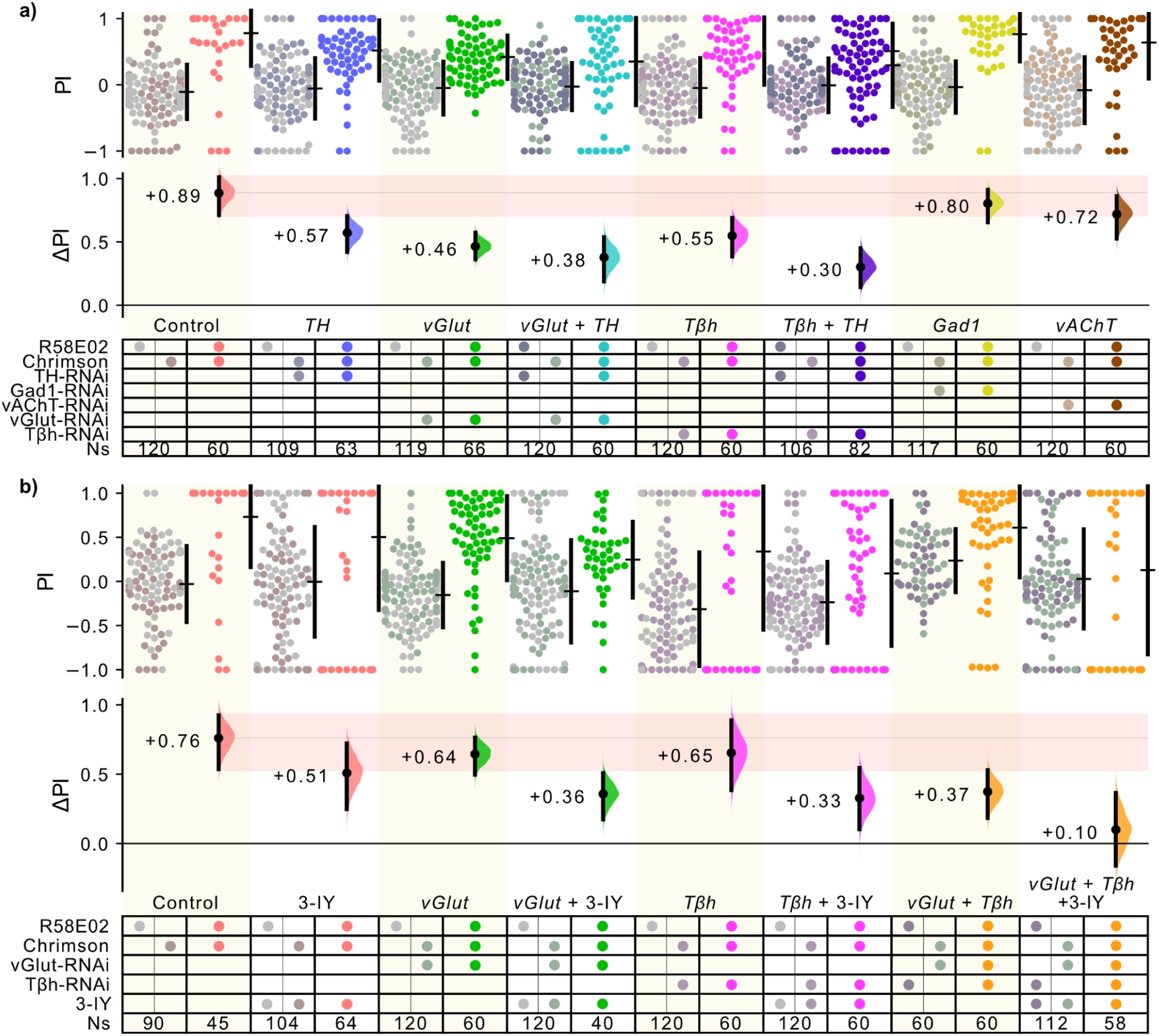
Large reductions in PAM valence require knockdown of multiple neurotransmitters. **a.** A knockdown screen for neurotransmitters that contribute to *R58E02* valence. *R58E02>Chr* flies were crossed with RNAi transgenes targeting factors required for five transmitters: *TH* (dopamine, replicating the prior experiment), *vGlut* (glutamate), GAD1 (GABA), vAchT (acetylcholine), and *TβH* (octopamine). The *vGlut* and *TβH* knockdowns showed a reduction in valence comparable to the TH knockdown. Simultaneous knockdown of TH with either *TβH* or vGlut resulted in a further reduction in the effect size. **b.** A combinatorial approach using 3-IY to systematically deplete dopamine along with RNAi-mediated knockdown of vGlut and *TβH* reveals a progressive reduction of valence with depletion of each neurotransmitter, with valence being virtually depleted when 3-IY, vGlut-RNAi and *TβH*-RNAi were simultaneously present (ΔPI = +0.10 [95%CI -0.16, +0.37]). Data for all panels can be found in the corresponding folder on the Zenodo data repository (https://doi.org/10.5281/zenodo.7747425).

Because the knockdowns of *TH, vGlut* and *Tꞵh* each resulted in a partial reduction of valence, we hypothesized that knocking down these genes in two-way combinations would elicit additive deficits. Indeed, the combined knockdown of *TH* and *vGlut* via the co-expression of both inhibitory RNAs produced a further reduction in valence, as did the simultaneous knockdown of *TH* and *Tꞵh* (**Figure 5a**); however, neither two-way combination succeeded in ablating valence completely. We thus hypothesized that all three neurotransmitters needed to be depleted to ablate valence. As the co-expression of all three RNAis with Chr proved to be technically prohibitive, we instead used the competitive inhibitor 3-IY to systematically reduce dopamine in conjunction with RNAi knockdowns. [Note that pseudo-replicate knockdowns of *vGlut* or *Tbh*, using different food, yielded somewhat different valence scores (**Figure 5a, b**)]. Simultaneous depletion of two transmitters—by combination of 3-IY with *vGlut-*RNAi, 3-IY and *Tꞵh*-RNAi, or both *vGlut* and *Tꞵh*-RNAis—showed further reductions in valence (**Figure 5b**). Finally, depletion of all three transmitters using 3-IY, *vGlut-*RNAi and *Tꞵh-*RNAi virtually ablated valence (**Figure 5b**). These results are consistent with the idea that *R58E02* valence is reliant on a combination of dopamine, glutamate and octopamine.

### PAM-β valence is partially dependent on dopamine

The opposing valence of *R58E02* and *R15A04* suggests valence heterogeneity in different subsets of PAM DANs. Indeed, numerous studies have found that valence-related behaviors like food-seeking, courtship, sleep, and appetitive memory are dependent on different MB sub-compartments and specific DAN subsets [28,34,43,52,54,60,82–85]. To identify the PAM types that drive valence, we screened 20 split-Gal4 lines [50,52] and identified several with valence, both negative and positive (**Figure 6a–b**). Of these lines, we focused on *MB213B,* as it had the strongest positive valence (**Figure 6b**). This line expresses in the PAM-β1 and PAM-β2 types (PAM4 and PAM10, respectively) with minor expression in the PAM11 (PAM-α1) cells [50] (**Figure 6c**, **Table S2, Movie S8**). As valence in *R58E02>Chr* flies is only partially dependent on dopamine, we interrogated *MB213B>Chr* dopamine dependency. Knocking down *TH* in PAM-β cells produced variable results (**Figure S5a-c)**; to resolve these replicate differences, we used meta-analytic averaging to calculate weighted mean-difference estimates at the 22 and 70 μW/mm^2^ light intensities [86]. This averaging showed that when compared to non-RNAi flies (**Figure 6d**), knockdown of *TH* elicited robust (though incomplete) valence reduction: –68% and –65%, respectively (**Figure 6e-f**). These effects are larger than the *TH-RNAi* knockdown effects for *R58E02>Chr*, which were –42% and –40% for the same light intensities (**Figure 5f**).

**Figure 6.**
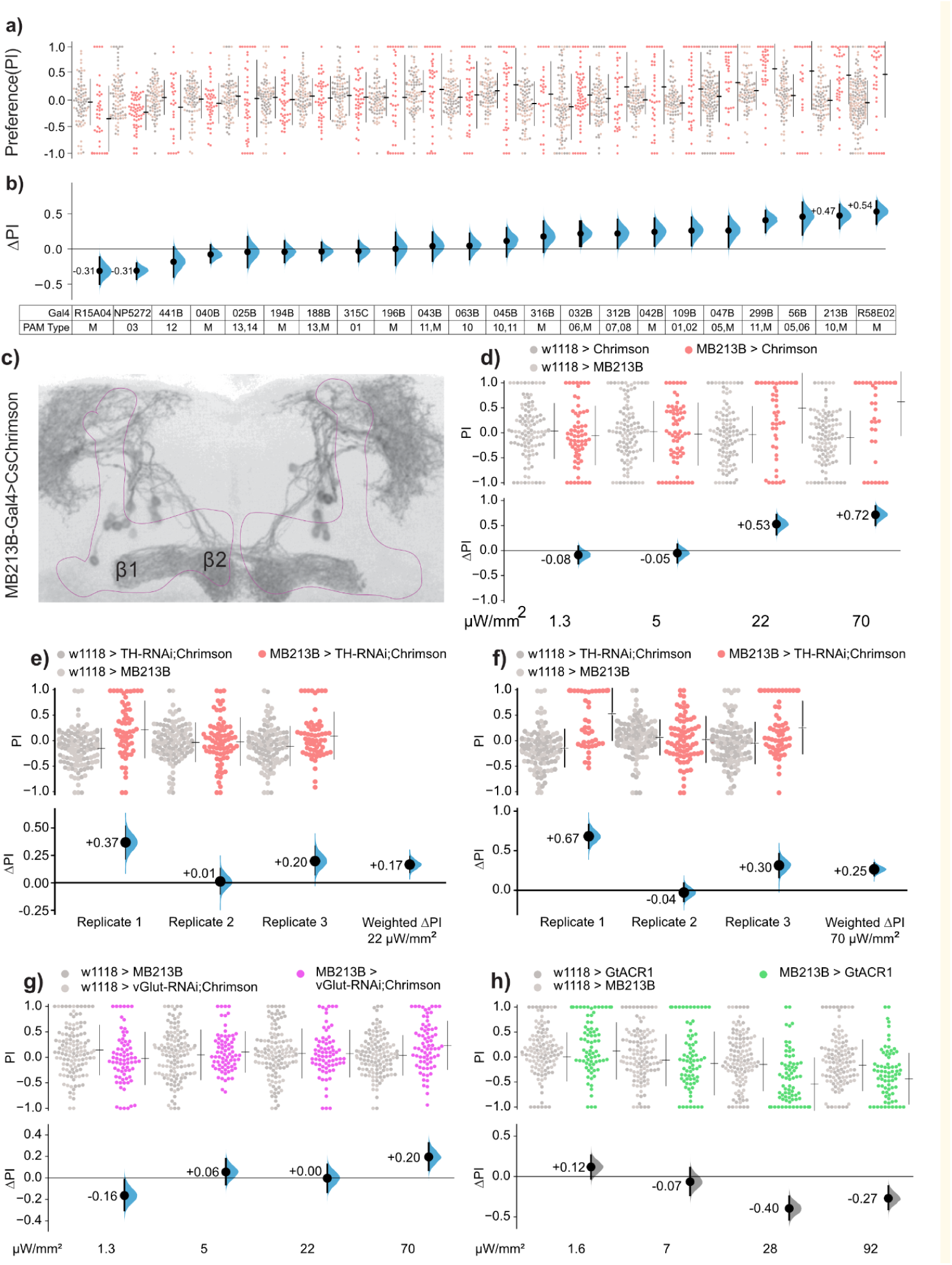
Valence mediated by dopaminergic PAM-β neurons is dependent on both dopamine and glutamate. **a-b.** An optogenetic activation screen of 22 PAM-DAN lines identified *MB213B* as the specifically expressing line with the strongest positive valence. Light preference was tested with 72 µW/mm^2^ red light. The table shows the PAM cell types in which each driver expresses; ‘M’ denotes multiple cell types; see **Table S2** for further details. **c.** The expression pattern of *Chr-YFP* with driver *MB213B*, showing projections to both zones of the β lobe (zones β1 and β2). **d.** Replication of the *MB213B>Chr* screen experiment confirmed that these flies are attracted to optogenetic light at the two highest intensities. **e-f.** Meta-analysis of three replicates of *MB213B > TH-RNAi; Chr* yielded weighted ΔPI values of +0.17 at 22 μW/mm^2^ and +0.25 at 70 μW/mm^2^ (orange curves). **g.** Expressing vGlut-RNAi with the MB213B driver similarly resulted in reduced (but not ablated) valence. **h.** *MB213>ACR1* flies avoided the green-illuminated area. Data for all panels can be found in the corresponding folder on the Zenodo data repository (https://doi.org/10.5281/zenodo.7747425).

### PAM-β valence is partially dependent on glutamate

PAM-β cells have been shown via single-cell RNA-seq to express several other neurotransmitter-related genes, including *vGlut*, *vAChT,* and *Gad1* [28]. As glutamate was required for *R58E02* valence (**Figure 5**), and is known as a co-transmitter in dopaminergic cells [87], we examined the effect of *vGlut* knockdown in the specific PAM-β driver. The *MB213B > vGlut-RNAi; Chr* flies displayed valence that was reduced but—like the *TH* knockdown—not completely abolished (**Figure 6g**). Thus, it appears that for valence mediated by the *MB213B* driver, glutamate and dopamine transmission each make partial contributions. These results show that as for *R58E02* valence, at least one specific PAM subset similarly drives acute valence using multiple transmitters.

### Silencing activity in β-lobe DANs drives a negative valence response

We next asked whether flies would respond to inhibition of the *MB213B* cells [57]. To drive inhibition, we expressed ACR1 in the *MB213B* cells [74,75]. The *MB213B>ACR1* flies avoided green light, i.e. showed negative valence. That ACR1 actuation of the PAM-β cells has a behavioral effect indicates that at least a subset of these cells were active in control flies during the experiment (**Figure 6h**).

### DANs and MBONs drive valence with different locomotion patterns

In olfactory learning, appetitive DANs are thought to be paired with aversive MBONs [88,89]. The concurrent release of dopamine with odor experience then causes weakening of the connections between the KCs and the MBONs, thus reducing the aversive response to the presented odor and driving approach [53,54,84,90]. Many MBONs themselves drive acute valence and receive synaptic input from DANs [51,52]. Considering that MBON types were shown to drive acute aversion, we hypothesized that optogenetic PAM activity might be inhibiting these aversive MBONs, and thereby rendering these output neurons the direct mediators of acute PAM valence.

To address this hypothesis, we screened a panel of MBON drivers for valence (**Figure 7a**, **S6a-b**). We found that one line, *VT999036*, had the strongest valence response. This line drives expression in MBONs that project to the γ lobe, termed MBON-γ1γ2 and MBON-γ4γ5 cells, also known as types MBON20 and MBON21, respectively [24,50] (**Figure 7b**, **S6e-f, Movie S6, Table S2**).

**Figure 7.**
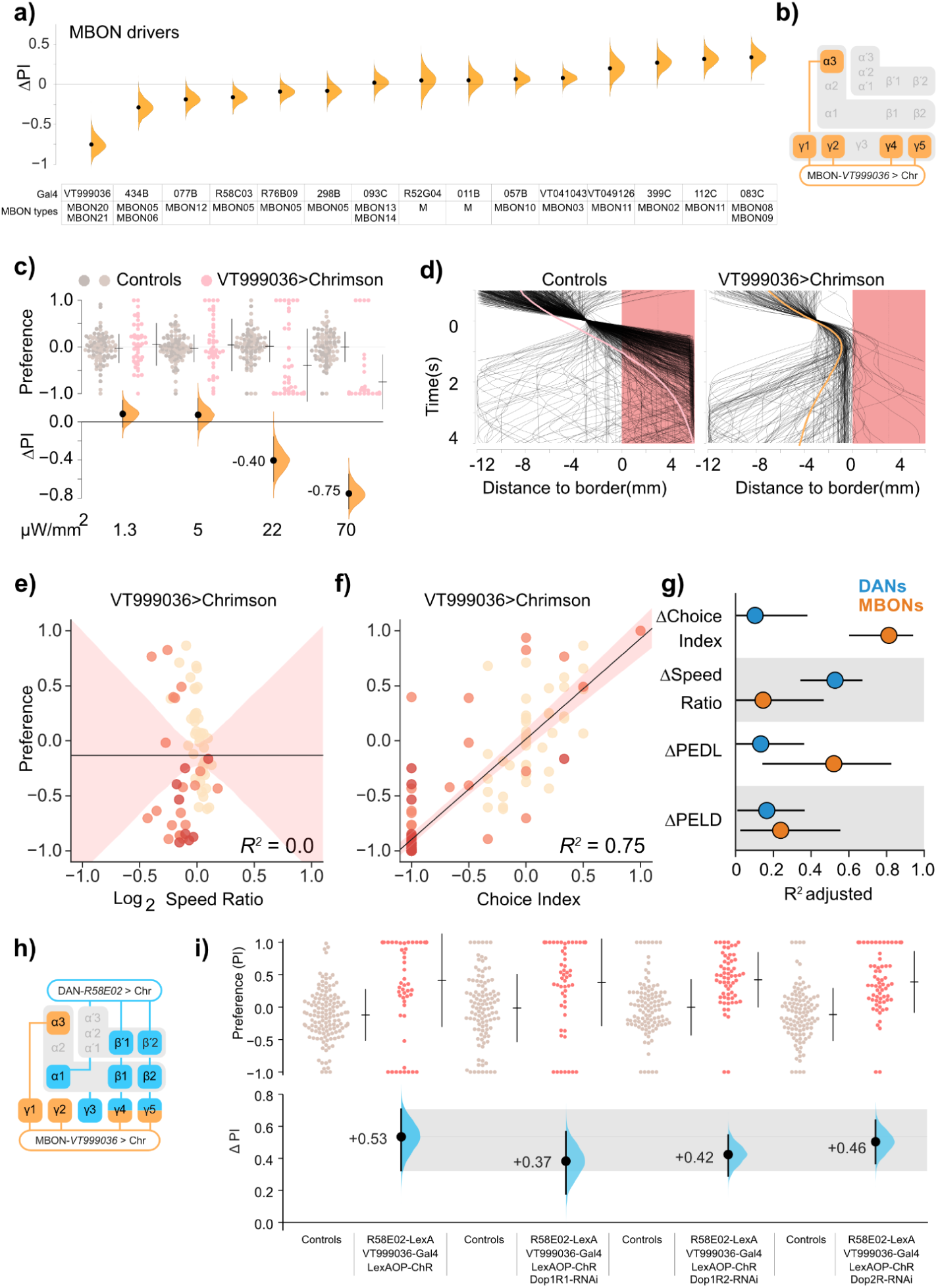
MBONs drive valence via choice effects, not speed effects. **a.** A screen of optogenetic Chr valence in 15 MBON-related lines (split-Gal4 and Gal4 drivers). Orange markers show the valence scores (black dots) and distributions (curves) of each cross, comparing test flies with controls. See **Table S1** for effect sizes. The matrix key shows driver identifiers in the top row, and MBON cell types in which each driver expresses; M denotes multiple cell types. See **Table S2** for further details. **b.** Schematic of *VT999036* projections to MBON synaptic-zone subsets. *VT999036* drives expression in two MBON types, MBON20 and MBON21 [50]. **c.** *VT999036>Chr* flies avoided opto-activation at the two highest illumination intensities (22 and 70 µW/mm^2^). The valence curve was produced using the same data from the screen summary. **d.** When *VT999036>Chr* flies move through the choicepoint, they tend to turn away from the light. **e.** In *VT999036>Chr* flies, a relationship between preference and speed ratios was absent. **f.** In *VT999036>Chr* flies, choice index and preference were related. **g.** Summary of regressions of DAN and MBON driver valence. Coefficients of determination for DAN lines (blue dots) and MBON lines (orange dots) are shown for four locomotor metrics as compared to valence (ΔPreference). The four metrics are Δchoice, Δspeed ratio, and the effect sizes of the dark-light and light-dark choice-point exit probabilities (ΔPEDL and ΔPELD, respectively). **h.** Combined schematic of combined *R58E02* and *VT999036* projections to the MB. The two expression patterns overlap in the γ4 and γ5 regions, corresponding to MBON21. **i.** Knocking down *Dop1R1, Dop1R2* and *Dop2R* in the MBONs of VT999036-Gal4 had modest effects on *R58E02>Chr* light preference. The gray ribbon indicates the control-valence 95% confidence interval. Data from 70 µW/mm^2^ red light. Data for all panels can be found in the corresponding folder on the Zenodo data repository (https://doi.org/10.5281/zenodo.7747425).

Activation of *VT999036* caused strong aversive valence, with flies turning away from the light at light–dark boundaries (**Figure 7c-d**, **Movie S7**). The ‘dark preference’ of *VT999036>Chr* flies had little correlation with light–dark speed differences, but was correlated with a choice index, i.e., fly trajectories at the boundary (**Figure 7f-g**). These locomotion patterns stand in contrast with *R58E02>Chr* and *R15A04>Chr* flies, in which valence correlated with optogenetic speed differences (**Figure 1g-h**, **S1c-f**).

These observations led us to ask whether the speed/choice dissociation observed in the *R58E02* and *VT999036* lines was part of a general trend for DAN and MBON lines. We analyzed metrics for all lines in the MBON and DAN screens (**Figure 6a-b**, **S4a-b, S6a-d**) and used them in regressions of the two screens’ valence and speed-ratio effect sizes. This analysis indicated that PAM-mediated valences were weakly determined by choice and strongly determined by speed differences (**Figure 7g**, **S7a-d**). By contrast, MBON-mediated valence was weakly associated with speed and more strongly determined by choice and choice-point exit probabilities (**Figure 7g**, **S7e-h**).

Because lines for both neuronal categories can drive attraction and avoidance, this difference is not easily explained by valence polarity. Nevertheless these data suggest that MBON and DAN activities have differential effects on two navigational properties: turning and speed.

### Dopamine-receptor knockdowns in strongly valent MBONs produce modest reductions in broad-PAM valence

As *VT999036* produced the strongest aversive valence in our MBON screen, we hypothesized that valence caused by broad PAM activation might be mediated via MBONs marked by *VT999036*, given the hypothesized pairing of appetitive DANs with aversive MBONs. The projections of *R58E02* and *VT999036* overlap in the γ4 and γ5 synaptic zones, corresponding to the MBON21 cells (**Figure 7h**).

Compared to the control effect size of ΔPI = +0.53 observed in *R58E02-LexA>LexAOp-Chrimson* flies, individual knockdowns of *Dop1R1* and *Dop1R2* in *VT999036* each produced a modest reduction in valence: approximately –30% and –21%, respectively (**Figure 7i**). Knocking down *Dop2R* resulted in only a –7% change in valence (**Figure 7i**). Notably, the summed effect of the *Dop1R1* and *Dop1R2* knockdowns (–51%) is comparable to the approximately –46% average reduction of valence observed earlier with dopamine-depleting interventions (**Figure 4i**). These results suggest that the two DopR1-like receptors in the MBON21 cells contribute to *R58E02* valence.

### Co-zonal PAM and MBON valences are not coherently related

Broad-PAM *R58E02* attraction is linked to *VT999036* aversion (as shown by targeted receptor knockdowns; **Figure 7i**), and both project to the γ4 and γ5 synaptic zones; these observations lend credence to the idea that acute positive DAN valence is mediated by inhibiting their postsynaptic aversive MBONs. To explore if this pattern extended beyond this instance, we explored whether PAM:MBON driver pairs with shared MB synaptic zones have an inverse valence relationship. Interestingly, the valence scores (ΔPIs) of co-zonal PAM and MBON did not show a consistent trend, contradicting the idea that co-zonal PAMs and MBONs have systematically opposing valence effects (**Figure S8**). For example, *MB213B* activity is consistently attractive, and these cells project to both of the two β zones (**Figure 6a-b, d**) [50]. Yet, two β-zone-projecting MBON drivers have variously positive and negative valence: *MB399C* (MBON02, zones β2 and β’2, ΔPI = +0.25) drives attraction; and *MB434B* (MBON05 and MBON06, zones ꞵ1 and γ4, ΔPI = –0.27) drives avoidance (**Figure S8**) [50]. It therefore seems that acute valence resulting from PAM stimulation does not necessarily arise from the inhibition of co-zonal aversive MBONs, or at least there is no clear evidence supporting a simple, general PAM:MBON valence relationship.

## Discussion

### Differences between appetitive learning and acute DAN valence

In this study we provide evidence that a dopaminergic system known to instruct learning also drives acute valence. The PAM DANs instruct appetitive odor learning [9,76], such that subsequent encounters with the same odor will elicit an increased approach behavior. Three features distinguish DAN-mediated olfactory learning from DAN-evoked acute valence. First, unlike classical Pavlovian learning—which requires an association between dopamine and a sensory stimulus—acute DAN valence occurs in an experimental environment that is otherwise largely featureless, such that the optogenetic illumination is the only salient sensory stimulus. Thus, the strong elicited valence observed is consistent with non-associative DAN functions that operate independently of external stimuli [45,54]. Second, while KCs are critical to DAN-mediated sensory associative learning [72,73], activated DAN acute valence has little-to-no reliance on KCs. Third, while learning has a strong dependency on *Dop1R1* receptors in the KCs [19,23,77]*, R58E02* PAM valence does not have critical dependencies on either dopamine-receptor function in KCs or PAM dopamine synthesis. Thus, DAN-dependent learning and DAN valence seem to be distinct processes that act through different circuits and signaling systems.

### Differences between DAN and MBON valence

Paired-posterior-lateral 1 (PPL1) DANs can drive optogenetic valence behavior [91]. Findings from our study exclude KCs as the downstream neurons through which PAM DANs affect valence. This finding suggests that MBONs are the possible mediators, as this class of neurons are major output cells of the MB [50,88,92,93] and receive synaptic input from DANs [91].

Under the assumption that DAN valence is mediated by MBONs, we explored some features of valent locomotion as driven by the two cell types. The DAN lines primarily influenced optogenetic light preference by affecting walking speed, while the MBONs, as previously shown [52], had their primary effect by changing trajectories at the light–dark interface, a trend that was independent of valence polarity. There are at least four possible explanations for this difference. First, some of the driver lines (e.g. *VT999036*) capture cells without MB projections, and these could (in some cases) contribute to valence. Second, DANs may have a symmetrical influence on walking, while MBONs have an asymmetrical effect on walking, perhaps via an algorithm similar to those in Braitenberg vehicles [94]. Third, assuming that DAN valence is mediated by MBON activity, it might be the case that MBONs have speed effects when quiescent, but drive turning when highly active, e.g., through specific downstream circuits with a distinct responsiveness to MBON activity. Fourth, the MBON screen might not have included the downstream cells responsible for DAN valence, i.e., other downstream circuits mediate DAN effects. On this fourth point: the drivers that express in co-zonal DAN and MBON types do not necessarily specify cellular subtypes that share synapses [61,95].

The specific function of DAN→MBON synapses in acute opto-valence remains unclear. Some MBON subtypes can modulate locomotor dynamics like walking speed and turning [52]; however, the extent to which DANs drive valent locomotion through MBONs is not known. Moreover, the idea that DAN valences are exclusively mediated by DAN→MBON synapses in the MB is likely to be incorrect. A connectomic analysis has shown that PAMs send axons to other neuropils [95], suggesting that at least some of the valence effects could be mediated through PAM signaling to non-MB areas.

### Appetitive DANs have non-associative functions

Overall, the speed/turning dissociation is hard to explain with current information. One possibility is that the visual associative pathway could send a sensory input to MBONs, thus providing a signal that involves the PAM DANs, but does not require the KCs. We do not consider this likely. While green light (like the light used to actuate ACR1) can indeed function as a visual conditioned stimulus, inhibition of KCs by *MB247-Gal4* abolishes visual learning [32]. In our hands, inhibition using the identical *MB247* driver left valence mostly intact, so the possibility of valence being equivalent to a conditioned visual response seems unlikely. Moreover, while olfactory-learning scores and *R58E02* optogenetic valence are of comparable magnitudes [96], visual-learning scores are typically <50% of this study’s valence effects [32,97]. These observations indicate that valence and visual learning are distinct.

The experimental results cast DAN-driven acute valence and olfactory sensory learning as separable processes, wherein DANs perform two distinct functions. In response to rewarding (or punishing) stimuli, PAM DANs seem to (1) write olfactory memories to Kandelian-type KC synapses for future reference, and (2) instruct immediate changes in locomotor behavior. Imaging of the MB has shown that DAN activity is closely connected with motor states and locomotion [45,54,98]. Inhibition experiments have revealed that DAN activities guide innate odor avoidance [99], and odor navigation [45]. Similarly, reward-related behaviors in mammals can be separated into consummatory, motivational, and learning components, of which the latter two are attributable to dopamine function [100]. Organizing parallel signals of a single reward circuit into distinct motivational and associative dopaminergic synapses could ensure coherence between valence (the present) and subsequent learned sensory responses (the future). From an evolutionary perspective, we speculate that the motor-function arm predates the associative arm [101], which was later inserted when the first sensory systems (taste and olfaction) developed.

### DAN valence relies on glutamate and octopamine

Knocking down *TH* function reduced *R58E02* valence by roughly half, suggesting the involvement of other neurotransmitters. Screening four other transmitter pathways implicated glutamate transport and octopamine synthesis; combined lesions showed that reduction of any two of *TH*, *Vglut*, and *Tbh* reduced *R58E02* valence by roughly two-thirds, while reduction of all three resulted in essentially no optogenetic valence. With the narrow PAM driver *MB213B*, knockdowns revealed that dopamine and glutamate were required by the PAM-β cells for normal valence. Previous single-cell RNA sequencing studies revealed the co-expression of *vGlut*, *Gad1*, *Tbh* and *vAChT* in subpopulations of dopaminergic neurons [28,102]. Together, these findings implicate dopamine, glutamate, and sometimes octopamine as co-transmitters in eliciting PAM-DAN optogenetic valence.

These findings are in agreement with a growing body of literature demonstrating the presence and utility of multiple transmitters in dopamine neurons and MB KCs. Recent work in PAMs has demonstrated roles for co-transmitters: memory updating requires the co-release of nitric oxide [28]; and appetitive memory formation is dampened by the co-release of GABA [103]. In PPL DANs, the dampening of aversive memory formation requires both GABA and glutamate [103]. Glutamate signaling from glia has also been implicated in aversive-memory formation, and octopamine-receptor knockdowns in KCs dampen aversive and appetitive-memory formation [104–107]. All of these prior studies focused on olfactory memory, whereas the experiments reported here demonstrate the non-associative functions of PAMs. More broadly, dopamine and glutamate co-transmission has been well-documented in mice, including in dopaminergic midbrain neurons, where dopamine and glutamate are released from distinct terminals [108–110]. Other neuromodulatory neurons also exhibit dual-transmission: some octopaminergic neurons in flies also transmit glutamate [111]. The use of multiple neurotransmitters thus seems to be a common mechanism by which neuromodulatory neurons exert their effects.

The molecular mechanisms by which dopamine, *Vglut* and *Tbh* act together to mediate valence are currently unknown. In *Drosophila*, synaptic vesicles at DAN terminals undergo hyper-acidification in response to neuronal activity; this process is driven by glutamate transport into these vesicles by *Vglut*, which in turn increases dopamine loading [87]. As a result, the effect of *Vglut* knockdown might be due to lower co-release of glutamate and/or reduced loading of dopamine into synaptic vesicles. As shown for some octopaminergic neurons that release octopamine and glutamate from separate termini [111], it is possible that dopamine and glutamate are also released from distinct PAM termini.

Normal *R58E02* valence also requires *Tbh*. While octopamine is known to be important in both appetitive and aversive memory [106], octopamine co-transmission from PAMs is novel. How octopamine-dependent, PAM-driven behaviors differ from DA- and glutamate-dependent behaviors are interesting topics for future studies.

### Technical note on interpreting knockdown data

Beyond the *TH-RNAi* immunostaining, we did not assess the molecular effects of the other RNAi manipulations, and it is likely that these RNAi transgenes produced partial knockdowns [78]. Previous work has shown that the *Vglut-RNAi* line used in this paper produces a ∼70% mRNA knockdown, whereas the *Gad1-RNAi* line produces an ∼82% protein knockdown [43,112]. The remaining lines (*Tbh-RNAi* and *vAChT-RNAi*) were sourced from the Transgenic RNAi Project: most of these lines produce knockdowns greater than 50% [113]. Nonetheless, that each of *vGlut-RNAi* and *Tbh-RNAi* produced a substantial reduction in *R58E02* valence demonstrates that the corresponding neurotransmitters have a role in *R58E02* valence. For *vAChT-RNAi* and *Gad1-RNAi*, neither produced appreciable changes to *R58E02* valence, but we cannot discern whether this was due to poor RNAi efficiency or that those transmitters have no role. Even though the individual knockdown effect sizes could be underestimated, nevertheless, the sum of the *TH*, *Vglut*, and *Tbh* knockdown effect sizes is approximately 100%, and the combination lesions indicate that reducing all three ablates valence almost completely (**Figure 3J**).

### Ongoing DAN activities shape behavior

That flies avoid silencing their PAM-β dopaminergic cells indicates that stimulus-independent DAN activities influence behavior. In the context of classical conditioning, dopamine is thought of as a transient, stimulus-evoked signal; this result indicates that some PAM neurons are active even in a chamber lacking odor, food, shock, or other salient stimuli. Physiological recordings show that, along with sucrose responses, PAM-γ activities correlate with motion and guide odor-tracking behavior, supporting the idea that PAM–DAN activities both respond to and steer locomotor behavior [45,54]. In our optogenetic preference screen, activity in various PAM DAN populations were rewarding, aversive or showed little preference effect, indicating that there is a diversity of functions between different DAN types, consistent with findings for learning and memory [9,11,14,20–28,76]. The bidirectionality of the attractive and aversive effects of increasing and decreasing activity in this dopaminergic system is similar to valence responses to activation and inhibition of dopaminergic cells in the mammalian ventral tegmental area [8,114,115] and is reminiscent of the increases and decreases in activity in that area that occur during positive and negative reward prediction errors, respectively [116]. Whole-brain imaging in the nematode has shown that global brain dynamics track closely with locomotion [117], suggesting that overarching brain function is to coordinate motor function. That the MB DANs drive preference-related locomotion suggest that their valence roles have two timescales: informing responses to future experiences, and steering current behavior.

### Discordance between PAM and MBON valence implies complex interactions

The knockdown of *Dop1R1* and *Dop1R2* in *VT999036* (MBON20, MBON21) produced reductions of valence similar to that of presynaptic dopamine depletion from the PAMs. Given that *VT999036* is highly aversive, it is tempting to hypothesize that PAM-mediated attractive valence acts by the inhibition of aversive MBONs, much in the way that memory is thought to be expressed [53,54,90]. Given the effects of the dopamine-receptor knockdowns, this paradigm might be the case for *VT999036*. When we examined the relationship between PAM and MBON valences across a range of drivers, however, it became clear that this was not a general pattern; that is to say, there was no consistent relationship between a PAM’s valence and the corresponding valence from its co-zonal MBONs.

The valence resulting from stimulation of a PAM subset seems to not arise from co-zonal MBONs, at least not exclusively. For memory, MBONs provide feedback to PAMs, not only via recurrent connections in the same zone but between zones as well [51,95,118–120]. Furthermore, MBONs do not have a one-to-one relationship with PAMs; there are more MBON cell types than PAM cell types and many MBONs sample information from multiple zones of the MB [50,51]. As such, when a population of PAMs is activated, it likely induces a complex network state that results in valence-related behaviors that may be mediated by an MBON outside of the PAM-targeted zone, or even by the overall effect of broader brain activity. Moreover, we cannot exclude the possibility that valence induced by stimulating different PAM subsets may work via diverse synaptic mechanisms, for example inhibiting aversive MBONs versus activating attractive MBONs. Further work is required to elucidate the circuit mechanisms required for PAM stimulation-driven valence, including imaging.

### Possible explanations for valence variability

Of this study’s many limitations, of note is the sometimes pronounced variability in results between experimental iterations. For example, *R58E02 > Chr* flies showed variable valence when tested by different experimenters at different times (**Figure 1e, 3g**). Similarly, *MB213B > Chr; TH-RNAi* flies showed variable valence (**Figure S5a-c**). One likely variance contributor is sampling error, a routine issue in behavioral data [121]. For several dopamine-depletion experiments, we adopted replication and meta-analysis to mitigate sampling error and estimate effect sizes with greater precision [96]. A second possibility is that neurotransmitter dependencies might vary between iterations due to uncontrolled changes during development. For example, the loss of dopamine may sometimes lead to developmental compensation, such as neurotransmitter switching or circuit adjustments. In mammals, transmitter switching in dopaminergic cells can occur as a result of stimuli such as odor or light stress [122–124]. In *Drosophila*, the expression of *vGlut* in DANs increases as dopamine is depleted either pharmacologically or due to aging [125]. It is possible that, as RNAi depletes TH, the PAM-β cells switch to glutamate as a substitute transmitter (but see the notes on effect-size surfeit above). A third possible explanation of the variability is that due to one or more uncontrolled variables, some neurons’ valence is susceptible to internal state. Imaging has shown that PAM-γ cells display activities that vary depending on state, such as starvation and walking [54,57]. If, for example, PAM-β baseline activity is high, it might be expected that the valence due to further activation will be small.

Conversely, in a low-baseline-activity state, stimulation of these cells would be expected to result in a large effect. Recent studies have described high variability between different optogenetic valence assays [62,65], suggesting that optogenetic valence is not generalizable between different behavioral tasks. As different sensory modalities and/or associative circuits may be employed for different tasks, this finding is not unexpected. Like all cognitive constructs, the broader property of so-called ‘valence’ will benefit from systematic investigations by multiple groups across behavioral paradigms.

### Valence modulation beyond PAM-DAN cells

The driver *R58E04-Gal4* mainly expresses in PAM–DAN cells, but it also shows expression in the optic lobes [10,25], raising the possibility that cells outside the PAM–DANs are responsible for some part of the valence. While this possibility could explain the residual valence after inducing dopamine lesions, we consider it to be unlikely. First, the cells in the optic lobe have been reported to be glia: “[*R58E02*] strongly labels the PAM cluster neurons and glial cells in the optic lobes with little expression elsewhere” [76]. Second, our screen of the PAM–DAN split-Gal4s shows that a number of them give positive valence, generalizing the phenomenon beyond the *R58E02* driver itself. For example, the *MB213B* split-Gal4 driver (which does not have any detectable expression outside of the PAM–DANs) has a positive valence that—like *R58E02*—is only partially reduced by dopamine knockdown. Third, the combination-knockdown experiments indicate that valence can be attributed to multiple transmitters, so an additional cause seems redundant. These observations support the idea that the valence observed is attributable to PAM–DAN activity.

### Questions beyond the study’s scope

Besides the questions we tackled in this study, there are other important areas worth considering. These include studying how the optogenetic activation of DANs and MBONs can give rise to distinct locomotor patterns. The role of co-transmission of different neurotransmitters in modulating these behavioral outcomes is another key aspect to explore. Furthermore, why certain DANs drive attraction while a subset of the larger DAN population induces aversion remains an intriguing question. Additionally, in the absence of olfactory stimuli, investigating the neurophysiological signal that dopamine acts upon to modulate walking speed is of interest. These questions extend beyond the current study and offer valuable avenues for future research.

Overall, our findings reveal that the PAM system, in parallel with its associative functions, can instruct acute valence behavior using distinct mechanisms. Associative learning shows a strong dependence on odor stimuli, KCs, and Dop1R1; by contrast, acute PAM valence does not have strong requirements for a salient olfactory stimulus, Orco-cell function, KC activity, or Dop1R1 reception in the KCs. The results reveal that the PAM neurons utilize multiple neurotransmitters such that dopamine is not acting alone, with glutamate and octopamine also having substantial roles in acute valence. The locomotor features of PAM valence seem distinct from MBON valence: while PAM valence is primarily mediated by a reduction in speed, MBON valence is mediated by a change in turning behavior. These findings provide insight into the diverse roles and mechanisms of PAM neurons in mediating both associative and non-associative valence responses, highlighting the complexity of this neuromodulatory circuit in regulating behavior.

## Methods

### Fly strains

Flies were cultured on a standard fly medium [126] at 25°C and 60% humidity in a 12 h light: 12 h dark cycle. Wild-type flies were a cantonized *w^1118^* line. The DAN and MBON split-Gal4 lines described in [50] were a gift from Gerry Rubin (Howard Hughes Medical Institute), except for VT041043-Gal4 [112] and VT49126-Gal4 [24], which were obtained from the Vienna Drosophila Resource Center (VDRC). *VT999036* was a gift from Barry Dickson (Howard Hughes Medical Institute). The Gal4 transgenic lines were obtained from the Bloomington Drosophila Stock Center (BDSC) and included: *R58E02-Gal4* [10], *R15A04-Gal4* [69], *20x-UAS-CsChrimson*[66], 13X-LexAOp2-GtACR1 [127], *MB247-Gal4*[128], *MB247-LexA* [129]*. R53C03-Gal4* [50], *R76B09-Gal4* [24], *R52G04-Gal4* [52].

*NP5272-Gal4* [16] were obtained from the Kyoto Stock Center (DGRC). The RNAi lines used were: Dop1R1 (KK 107058), Dop1R2 (KK 105324), Dop2R (GD 11471), vGlut (KK104324), and TH (KK 108879), obtained from VDRC; as well as Gad1 (BDSC_51794), Tꞵh (BDSC_76062), and vAChT (BDSC_80435) from the Transgenic RNAi Project [113]. Supporting table in **Table S1** provides detailed descriptions of genotypes shown in each figure.

### Transgenic animal preparation

*Gal4*, *UAS-CsChrimson*, and *UAS-ACR1* crosses were maintained at 25°C and 60% humidity, in darkness. Groups of 25 newly eclosed flies were separated into vials for 2–3 days (in the dark at 25°C) before behavioral phenotyping. Control flies were generated by crossing the driver or responder line with a wild-type *w^1118^* strain (originally bought from VDRC), and raising the progeny under identical regimes to those used for the test flies. A stock solution of all-*trans*-retinal was prepared in 95% ethanol (w/v) and mixed with warm, liquefied fly food. Each vial was covered with aluminum foil and incubated at 25°C in the dark. Before optogenetic experiments, 3–5 day-old male flies were fed 0.5 mM all-*trans*-retinal (Sigma) for 2–3 days at 25°C in the dark.

### Drug treatment

Male flies (3–5 days old) were placed on 1% agar containing 5% sucrose, 10 mg/mL 3-iodo-L-tyrosine (3-IY, Sigma), and 0.5 mM all-trans-retinal for 2–3 days at 25°C in the dark prior to behavioral testing. Control flies were fed on the same food but with 3-IY omitted.

### Immunohistochemistry

Immunohistochemistry was performed as previously described [75]. Briefly, brains were dissected in phosphate buffered saline (PBS) and fixed in PBS with 4% paraformaldehyde (Electron Microscopy Sciences) for 20 min. Samples were washed three times with PBT (PBS + 1% Triton X-100) and blocked with 5% normal goat serum for 1 h. Samples were then incubated with primary antibodies overnight at 4°C. After three additional washes with PBT, samples were incubated with secondary antibodies overnight at 4°C. The following primary and secondary antibodies were used: mouse α-DLG1 (4F3 α-DISCS LARGE 1, Developmental Studies Hybridoma Bank, 1:200 dilution), rabbit α-TH (AB152, Millipore, 1:200 dilution), chicken α-GFP (Abcam, ab13970, 1:1000), Alexa Fluor 488 rabbit α-GFP-IgG (A-21311, Invitrogens, 1:200), Alexa Fluor 568 goat anti-mouse (A-11004, Invitrogen, 1:200), Alexa Fluor goat anti-chicken 488 (A-11039, Invitrogen, 1:200), Alexa Fluor goat anti-rabbit 568 (A-11036, Invitrogen, 1:200).

### Confocal laser microscopy and neuroanatomy

Confocal images were acquired under a Zeiss LSM 710 microscope at a z-step of 0.5 μm using 20×, 40×, or 63× objectives. Images were analyzed using ImageJ software. Black and white images are a maximum projection intensity (MIP) of the green channel. The stacks were visualized and analyzed with the FIJI distribution (**Error! Hyperlink reference not valid.** of ImageJ (NIH). Outlines of α-Dlg1 expression in the mushroom body were traced with Adobe Illustrator. Projection patterns and zonal identity were assigned as previously described [50]. When not verified by microscopy, cell types and projection patterns were classified by review of published reports (**Table S2**) [24,25,50,60,76,112].

### Optogenetic response assay

Behavior experiments were performed as previously described [75]. Each behavioral arena was cut with 55 × 4 mm stadium/discorectangle geometry; 15 such arenas were cut from 1.5 mm-thick transparent acrylic. During the behavioral assay, arenas were covered with a transparent acrylic lid. As previously described [3], flies were anesthetized on ice before loading into each chamber in the dark. The arena multiplex was kept under infrared (IR) light at 25°C for 2–3 min before starting the assay. Flies were aroused by shaking the arenas just before starting the experiment. All behaviors were recorded under IR light. The multiplex was illuminated with red or green light from a mini-projector positioned above the arena (Optoma ML750). For *CsChrimson* experiments, flies were illuminated with four red-light intensities: 1.3, 5, 22, and 70 µW/mm^2^. For *ACR1* experiments, the flies were illuminated with four green-light intensities: 1.6, 7, 28, and 92 µW/mm^2^. The colored light intensity was varied by changing the level of the respective RGB component of the projected color. For each experiment, the arenas were illuminated for 60 s with equal-sized quadrants to produce a banded light-dark-light-dark pattern.

### Video tracking

The behavior arena was imaged with a monochrome camera (Guppy-046 B, Allied Vision) with two IR longpass filters in series (IR Filter IR850, Green.L). Videos were processed in real time with CRITTA software written in LabView [75]. The x-y coordinates of each fly’s head were individually tracked (at 25 frames per second) using CRITTA’s tracking feature. CRITTA was also used to control the timing, hue and intensity of the illumination, and to count the number of flies in each quadrant for each video frame. The light borders were identified and calibrated using a function of the CRITTA plugin, which illuminates quadrants at low intensity and captures an image of the arenas (with camera IR filters removed). The plugin software calculates the horizontal intensity profile of each arena and finds the center of each light–dark boundary using an edge-detection algorithm. The light–border drift between presented experiments was 330 µm (95CI 230 µm; 430 µm). Between the light and dark regions was a light gradient that was a mean 670 µm wide with a range of 420 to 1040 µm. This gradient was measured from 45 images of boundaries from 15 chambers, and scored as all the pixels falling between high (light-on) and low-intensity light regions.

### Olfactory conditioning

Conditioning was performed as previously described [75,130]. Briefly, each behavior chamber was 50 mm long, 5 mm wide, and 1.3 mm high; the floor and ceiling of each chamber were composed of transparent shock boards made from indium tin oxide electrodes printed on glass (Visiontek UK). Odorized air was pumped into the ends of each arena at 500 mL/min. The odors were 3-methylcyclohexanol (MCH) at 9 parts per million (ppm) and 3-octanol (OCT) at 6 ppm, as measured with a photoionization detector (RAE systems, ppbRAE3000). The air exited the chamber via two vents located in the middle, creating two odor partitions in the conditioning area. Each experiment was performed with 4–6 flies. For opto-conditioning, flies were presented with either OCT or MCH odor paired with green light (515 nm, 28 µW/mm^2^), followed by another odor without visible light (IR light only). During shock conditioning, the presentation of either OCT or MCH was coupled with 12 electric shocks of 1 s duration at 60V [14]. Conditioned-odor preference (memory) was tested by the presentation of both odors, one from each side. For each of the two odors, a half performance index (PI) was calculated according to the fly position coordinates during the last 30 s of each testing phase; for each iteration, data from odor pairs were averaged to obtain a full PI [131].

### Preference and speed analysis

Custom Python scripts were used for data processing, analysis and visualization. The scripts integrated several routines from NumPy, pandas, matplotlib, and seaborn. For every fly, the x-y coordinates of the head location (recorded at 25 frames per second) underwent rolling-window smoothing, using a centered 1 s-wide triangular window. The following metrics were obtained (for every fly) for the last 30 s of each test session:

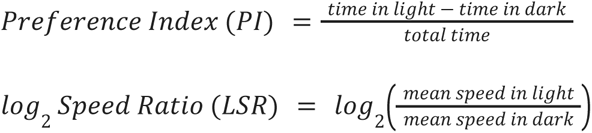

While all flies could be assigned a PI, the log_2_ Speed Ratio (LSR) could be computed only for flies that moved in both light and dark regions during the illumination epoch. Flies that remained stationary for the entire illumination epoch, or remained in only the light or dark zones, were excluded from the speed ratio calculation. Flies that started and remained in either the dark or light zone throughout an epoch, but still moved within the zone, were assigned an extreme PI (–1.0 or +1.0, respectively).

### Choice-zone trajectory analysis

A choice trajectory was defined as any transit in and out of a choice zone defined to extend 3 mm in either direction from all three light borders.

Trajectories were identified for every fly that approached the choice zone, and the following metrics were computed:

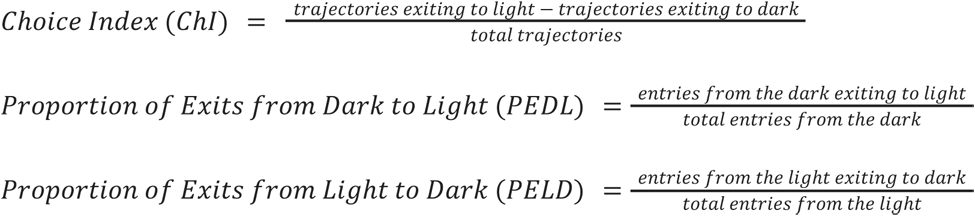

The above three metrics were computed for all flies that entered a choice zone at least once during the illumination epoch. Flies that did not make such a crossing during the epoch (i.e. remained on one side for the epoch duration) were necessarily excluded from boundary trajectory analysis. Note that as flies could enter a choice zone without ever subsequently crossing from dark to light (or vice versa), not all flies with a choice index could also be assigned a speed ratio; this would include flies that consistently made choice-zone reversals without crossing a light–dark boundary. Thus, the choice-zone analysis necessarily excluded flies that never crossed a light–dark boundary.

### Effect-size regression

To compare the valence effect size (Δ Preference or ΔPI) with locomotion effect sizes, Δ values were calculated for four locomotion metrics.

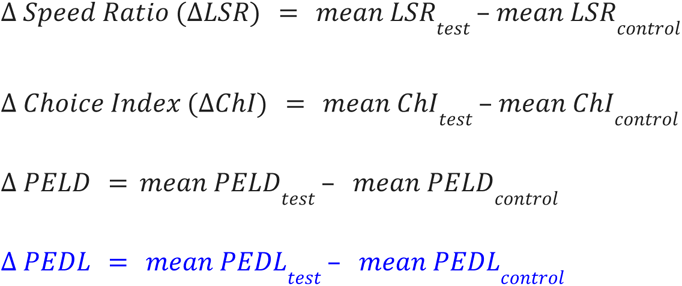

Each Δ value was calculated for the two highest illumination intensity epochs (22 and 70 µW/mm^2^) for all PAM or MBON lines. As the locomotion metrics require that a fly crosses the light–dark boundary at least once, some flies were necessarily censored from this analysis.

Each set of effect sizes was subjected to regression against the corresponding Δ Preference values. Regression was performed with the linear least-squares method of the SciPy library. For both mean differences and coefficients of determination (R^2^), distributions and 95% confidence intervals were obtained from 3,000 resamples, using bias correction and acceleration [132] with the scikits.bootstrap package.

### Meta-analysis of dopamine-depleting interventions

The dopamine loss-of-function experiments with *R58E02>Chr* were established using 3-iodo-tyrosine, *TH-RNAi*, or *TH-RNAi* with *Dicer2* at several light intensities, resulting in a total of 15 valence experiments. The effect of reducing dopamine function in the *R58E02* cells was estimated as a percentage of wild-type behavior [96,133]. Using data from three replicates of the *R58E02>Chr* experiment, a mean ΔPI (controls) valence was first calculated via simple averaging for the 5, 22 and 70 μW/mm^2^ light conditions. The ΔPIs for each dopamine-depleting intervention were then expressed as a percentage of the control value [96] for each of three light intensities .

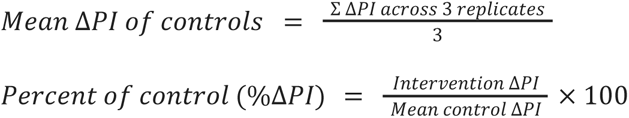

The data used for calculating the mean ΔPI (controls) value are presented in **Figures 3g**, **S3a** and **S3b**. The data used for calculating % of control for each intervention are presented in figures: **S3g** (3-iodo-tyrosine); **3h**, **S3c** and **S3d** (*TH-RNAi*); **S3e** (*TH-RNAi* with *Dicer2*).

### Meta-analysis of experimental replicates

For replicates of the *MB213> TH-RNAi; Chrimson* experiment, an inverse-variance meta-analysis was performed with a fixed-effects model. A weighted effect size was calculated as follows:

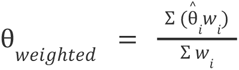

Where:

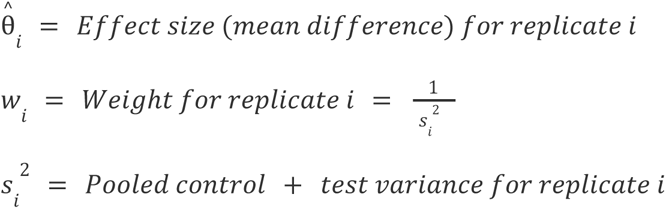

### Correlation analysis for PAM:MBON preference-index effect sizes

PAM:MBON driver pairs for analysis in **Figure S8** were identified on the basis of both drivers in each pair having major anatomical staining in the same MB zone. For lines from the FlyLight split-Gal4 collection, major staining was defined as a staining intensity of 3 or greater, as defined in a previous report [50]. For other, non-split Gal4 driver lines, the presence of major zonal projections was derived from other previously published reports (**Table S2**).

Regression was performed with the linear least-squares method of the SciPy library.

### Statistics

Effect sizes were used for data analysis and interpretation. Summary measures for each group were plotted as a vertical gapped line: the ends of the line correspond to the ±standard deviations of the group, and the mean itself plotted as a gap. Effect sizes were reported for each driver line as mean differences between controls and test animals for all the behavioral metrics [134]. Two controls (driver and responder) were grouped together and the averaged mean of the two controls was used to calculate the mean difference between control and test flies. In text form, the mean differences and their 95% confidence intervals are presented as “mean [95CI lower bound, upper bound].” The mean differences are depicted as black dots, with the 95% confidence interval indicated by error bars. Where possible, each error bar is accompanied with a filled curve displaying the distribution of mean differences, as calculated by bootstrap resampling. Bootstrapped distributions are robust for non-normal data [135].

*P* values were calculated by the Mann-Whitney rank method in SciPy and presented *pro forma* only: following best practice, no significance tests were conducted [134]. The behavioral sample sizes (typically *N =* 45, 45, 45) had a power of >0.8, assuming α = 0.05 and an effect size of Hedges’ *g* = 0.6 SD. All error bars for mean differences represent the 95% confidence intervals.

Genotypes and statistics for control and test flies for each panel are provided in the supporting table (**Table S1**), as are details of the mega-analysis of mutant effect sizes (**Figure 4g**) incorporating data from across the study.

### Data availability

All data generated for this paper are available for download from Zenodo (https://doi.org/10.5281/zenodo.7747425).

### Code availability

All code used for analysis of data is available for download from Zenodo (https://doi.org/10.5281/zenodo.7747425).

## Acknowledgments

The authors would like to thank Gerald Rubin (Howard Hughes Medical Institute) for the DAN and MBON split-Gal4 lines, Barry Dickson (Howard Hughes Medical Institute) for providing the *VT999036* flies. The authors thank Aldrich Hezekiah for illustrations. The authors thank Dr. Jessica Edwards of Insight Editing London for critical review of the manuscript before submission.

## Author Contributions

Conceptualization: FM and ACC; Experiment design: FM, YM and ACC; Methodology: FM, JCS and ACC; Software: JH (Python), and JCS (CRITTA, LabView); Data Analysis: JH (Python), FM (Python), YM (Python) and JCS (LabView); Investigation: FM and YM (genetics, fly husbandry, behavior, immunohistochemistry, and microscopy), XYZ (behavior, brain dissection, immunohistochemistry, microscopy), SO (memory); Resources: JCS (instrumentation); Writing – Original Draft: FM; Writing – Revision: FM, YM, and ACC; Visualization: FM, YM, JH, ACC; Supervision: ACC; Project Administration: ACC; Funding Acquisition: ACC.

## Legends for Supporting Information

## Supplementary Figures

**Figure S1. Dopaminergic-line optogenetic preference is related to changes in walking speed.**

**a-b.** Maximum-intensity projections of genetically-stained mushroom bodies show innervation by *R58E02* and *R15A04* in various MB neuropil zones. Dark pixels indicate α-GFP signal from the GFP tag on ACR1. *R58E02* has stained fibers across nearly all the horizontal lobes. *R15A04* sends fibers to four zones, α1, β2, β’1, and γ5. The MB lobes (purple line) were outlined using DLG-1 counterstain, not shown here (see Methods for details)—scale bar: 20 μm. See also Movies S1 and S3.

**c.** Scatter plots and linear regression of *R58E02>Chr* preference indices (PIs) and their light/dark log_2_ speed ratios. Each point indicates metrics for a single fly. The two metrics are negatively correlated: R^2^ = 0.44 (95 CI, 0.24, 0.62, *P* = 2.9 × 10^-12^). *N* = 86. Each point indicates the log_2_ speed ratio and PI of a single fly. Flies were assayed at 1.3, 5, 22, and 70 µW/mm^2^.

**d.** There is an inverse relationship between *R15A04>Chr* PIs and speed ratios; R^2^ = 0.45 (95 CI, 0.3, 0.6, *P* = 1.2 × 10^-20^).

**e-f.** No correlation between PI and choice index was observed in *R58E02>Chr (*R^2^ = 0.01 (95 CI, 0.00, 0.07, *P* = 3.4 × 10^-1^) *and R515A04>Chr (*R^2^ = 0.04 (95 CI, 0.00, 0.05, *P* = 3.1 × 10^-1^)flies; . Each point indicates metrics for a single fly. Flies were screened at 1.3, 5, 22, and 70 µW/mm^2^.

**Figure S2. Optogenetic PAM DAN valence does not require olfactory-system function.**

**a.** *R58E02-LexA>LexAop-Chr* flies are attracted to green light. The previous green-light optogenetic activation experiment (Fig. 3a) was replicated using the *R58E02-LexA* driver (instead of *R58E02-Gal4*) and reproduced the attraction phenotype.

**b.** In *R58E02>Chr/MB247>ACR1* flies, attraction to DAN self-activation is largely unaffected by simultaneous KC opto-inhibition (Fig. 3b). The green-light dual optogenetic experiment was replicated with *R58E02-LexA* and *MB247-Gal4* drivers; it reproduced the outcome observed in the previous experiment: valence is unaffected by inhibition of the *MB247* cells.

**c.** *Orco>ACR1* flies display moderate attraction to green light at two intermediate intensities (7 and 28 µW/mm^2^), but not at 92 µW/mm^2^.

**Figure S3. Drug-induced and enhanced RNAi-mediated depletion of dopamine results in partially reduced valence.**

**a-b.** Additional replicates of *R58E02>Chrimson* used in Fig. 4i in the main text (see Methods for details). Data from Fig. S3e is also used in column 1 of main text Fig. 5a.

**c-d.** Additional replicates of *R58E02>TH-RNAi;Chrimson* used in main text Fig. 4i in the main text (see Methods for details).

**e.** Enhancement of RNAi-mediated TH knockdown via the overexpression of *Dicer2* resulted in valence at 22 μW/mm^2^ of ΔPI = +0.26 [95CI +0.11, +0.39]; valence at 70 μW/mm^2^ was ΔPI = +0.53 [95CI +0.39, +0.66].

**f-g.** Depletion of dopamine *via* feeding with 3-iodotyrosine (10 mg/mL 3-IY) resulted in a moderate reduction of valence. At 70 μW/mm^2^, *R58E02>Chr* flies that were not fed 3-IY displayed valence of ΔPI = +0.76 [95CI +0.54, +0.92], whereas flies that were fed 3-IY displayed valence of ΔPI = +0.51 [95CI +0.25, +0.72]. The data from Fig. S3f–g is also used in columns 1–2 of the main text Fig. 5b.

**Figure S4. Corresponding log2 speed ratios for PAM-DAN valence screen.**

**a-b.** Δ Log_2_ speed ratios for the 22 PAM-DAN lines in our valence screen (**Fig. 6a-b**). Overall, lines with negative valence had positive speed ratios, while lines with positive valence had negative speed ratios. See **Table S1** for effect sizes. The table shows the PAM cell types where each driver expresses most strongly, and M denotes multiple cell types (DOI: 10.5281/zenodo.7239106). See **Table S2** for further details on driver expression.

**Figure S5. TH knockdown in PAM-β cells led to variable reductions in effect size.**

**a-c.** Replicates of MB213B > TH-RNAi; Chrimson experiments showed varying effect sizes, necessitating meta-analysis. These panels correspond to Replicates 1, 2 and 3, respectively, in **Fig. 6e-f**.

**Figure S6. In MBONs output synaptic zones, dopamine signaling is dispensable.**

**a-b.** An optogenetic valence screen for affective output zones using 15 MBONs related to split-Gal4 and Gal4 drivers. The data in panel b is a duplicate of Fig. 7a.

**c-d.** Δ Log_2_ speed ratios for the 15 MBON lines in the optogenetic valence screen. See Table S1 for effect sizes. The matrix key below indicates the drivers used and the corresponding MBON cell types in which they express; see Table S2 for details.

**e-f.** Immunostaining of YFP in MBONs labeled by VT999036-Gal4. See also **Movie S6**.

**Figure S7. Regression analyses of DAN and MBON drivers’ behavior metrics.**

**a-d.** Scatter plots relating valence with Δchoice (ΔChI), Δspeed ratio, Δproportion of dark→light exits (ΔPEDL) and Δproportion of dark→light exits (ΔPELD) in the DAN lines.

**e-h.** Scatter plots relating valence to Δchoice, Δspeed ratio, Δproportion of dark→light exits (ΔPEDL) and Δproportion of dark→light exits (ΔPELD) for the MBON lines from Fig. 7a and S7a. The data are obtained from the two bright illumination intensities (22 and 70 µW/mm^2^).

**Figure S8. No correlation between valence for co-zonal PAM and MBON activation.**

Valence scores from the PAM and MBON screens were compared for driver lines with similar projection patterns in the MB lobes. Each dot indicates the two valence scores from a pair of PAM:MBON drivers with major staining in identical neuropil zones. Error bars represent the confidence intervals of the valence (ΔPI) scores. For each pair, the PAM driver is denoted by color, while the MBON driver is denoted by marker style.

## Supporting Information: Tables

**Table S1. Detailed genotypes and effect sizes**

Genotypes and effect sizes of experiments in each figure. Semicolons indicate different chromosomes, commas indicate different transgenes. Compound genotypes are stated for split-Gal4 drivers, with the common line designation (e.g. MB213B) in parentheses.

**Table S2. PAM and MBON types**

A catalog of published information about the drivers used in the PAM (Fig. 5a-b) and MBON (Fig. 4a, S4a) screens, the cell types they express in, and the lobes of the MB in which each cell type’s terminals can be found.

## Supporting Information: Movies

**Movie S1. Z-stack for *R58E02>ACR1* immunostaining.**

Full confocal stack of the PAM cluster in a representative *R58E02>ACR1* fly brain, corresponding to Fig. S1a. Green: YFP; Magenta: ɑ-Dlg1; taken using 20× objective lens.

**Movie S2. *R58E02>Chr* flies walk slower in light**

Video of a 91-s-duration OSAR experiment, showing the dark phase and the first light epoch. *R58E02>Chr* flies slow or stop upon entering the light. The video is sped up to 1.33× realtime. Optogenetic light was turned on at t = 22 s and off at t= 106 s (video time). The light-border pattern was superimposed on the video (see Methods for details).

**Movie S3. Z-stack for *R15E02>ACR1* immunostaining.**

Full confocal stack of a subset of PAM neurons in a representative *R15E02>ACR1* fly brain, corresponding to Fig. S1b. Green: YFP; Red: ɑ-Dlg1; taken using 20× objective lens.

**Movie S4. Z-stack for *R58E02>Chr* immunostaining.**

Full confocal stack of the PAM cluster in a representative *R58E02>Chr* fly brain, corresponding to **Fig. 3e**. Green: ɑ-YFP; Red: ɑ-TH; taken using 63× objective lens. High colocalization between the YFP and TH signals is evident.

**Movie S5. Z-stack for *R58E02>TH-RNAi;Chr* immunostaining.**

Full confocal stack of the PAM cluster in a representative *R58E02>TH-RNAi;Chr* fly brain, corresponding to **Fig. 3f**. Green: ɑ-YFP; Red: ɑ-TH; taken using 63× objective lens. The colocalization between YFP and TH signals is reduced.

**Movie S6. Z-stack for *VT999036>ACR1* immunostaining.**

Full confocal stack of a subset of MBON neurons in a representative *VT999036>ACR1* fly brain, corresponding to Fig. S1b. Green: YFP; Magenta: ɑ-DLG; taken using 20× objective lens.

**Movie S7. *VT999036>Chr* flies avoid light**

*VT999036>Chr* flies avoid light using reversal and turning maneuvers. The timing is identical to that in Movie S2.

**Movie S8. Z-stack for *MB213B>Chr* immunostaining**

Full confocal stack of a representative *MB213B>Chr* fly brain, corresponding to **Fig. 5c**. Green: YFP; Magenta: ɑ-DLG; taken using 20× objective lens.

## References

1. Anderson DJ, Adolphs R. A framework for studying emotions across species. Cell. 2014;157: 187–200.

2. Darwin C. The Expression of the Emotions in Man and Animals. Project Gutenberg; 1899.

3. Mohammad F, Aryal S, Ho J, Stewart JC, Norman NA, Tan TL, et al. Ancient Anxiety Pathways Influence Drosophila Defense Behaviors. Curr Biol. 2016;26: 981–986.

4. Bargmann CI. Beyond the connectome: how neuromodulators shape neural circuits. Bioessays. 2012;34: 458–465.

5. Marder E. Neuromodulation of neuronal circuits: back to the future. Neuron. 2012;76: 1–11.

6. Schultz W. Multiple dopamine functions at different time courses. Annu Rev Neurosci. 2007;30: 259–288.

7. Bromberg-Martin ES, Matsumoto M, Hikosaka O. Dopamine in motivational control: rewarding, aversive, and alerting. Neuron. 2010;68: 815–834.

8. Namburi P, Al-Hasani R, Calhoon GG, Bruchas MR, Tye KM. Architectural Representation of Valence in the Limbic System. Neuropsychopharmacology. 2016;41: 1697–1715.

9. Burke CJ, Huetteroth W, Owald D, Perisse E, Krashes MJ, Das G, et al. Layered reward signalling through octopamine and dopamine in Drosophila. Nature. 2012;492: 433–437.

10. Liu C, Plaçais P-Y, Yamagata N, Pfeiffer BD, Aso Y, Friedrich AB, et al. A subset of dopamine neurons signals reward for odour memory in Drosophila. Nature. 2012;488: 512–516.

11. Lin S, Owald D, Chandra V, Talbot C, Huetteroth W, Waddell S. Neural correlates of water reward in thirsty Drosophila. Nat Neurosci. 2014;17: 1536–1542.

12. Han KA, Millar NS, Grotewiel MS, Davis RL. DAMB, a novel dopamine receptor expressed specifically in Drosophila mushroom bodies. Neuron. 1996;16: 1127–1135.

13. Kim Y-C, Lee H-G, Han K-A. D1 dopamine receptor dDA1 is required in the mushroom body neurons for aversive and appetitive learning in Drosophila. J Neurosci. 2007;27: 7640–7647.

14. Claridge-Chang A, Roorda RD, Vrontou E, Sjulson L, Li H, Hirsh J, et al. Writing memories with light-addressable reinforcement circuitry. Cell. 2009;139: 405–415.

15. Schroll C, Riemensperger T, Bucher D, Ehmer J, Völler T, Erbguth K, et al. Light-Induced Activation of Distinct Modulatory Neurons Triggers Appetitive or Aversive Learning in Drosophila Larvae. Curr Biol. 2006;16: 1741–1747.

16. Aso Y, Herb A, Ogueta M, Siwanowicz I, Templier T, Friedrich AB, et al. Three dopamine pathways induce aversive odor memories with different stability. PLoS Genet. 2012;8: e1002768.

17. Busto GU, Cervantes-Sandoval I, Davis RL. Olfactory learning in Drosophila. Physiology . 2010;25: 338–346.

18. Schwaerzel M, Monastirioti M, Scholz H, Friggi-Grelin F, Birman S, Heisenberg M. Dopamine and octopamine differentiate between aversive and appetitive olfactory memories in Drosophila. J Neurosci. 2003;23: 10495–10502.

19. Kim Y-C, Lee H-G, Han K-A. D1 dopamine receptor dDA1 is required in the mushroom body neurons for aversive and appetitive learning in Drosophila. J Neurosci. 2007;27: 7640–7647.

20. Krashes MJ, DasGupta S, Vreede A, White B, Armstrong JD, Waddell S. A neural circuit mechanism integrating motivational state with memory expression in Drosophila. Cell. 2009;139: 416–427.

21. Aso Y, Siwanowicz I, Bräcker L, Ito K, Kitamoto T, Tanimoto H. Specific dopaminergic neurons for the formation of labile aversive memory. Curr Biol. 2010;20: 1445–1451.

22. Aso Y, Herb A, Ogueta M, Siwanowicz I, Templier T, Friedrich AB, et al. Three Dopamine Pathways Induce Aversive Odor Memories with Different Stability. Rulifson E, editor. 2012. doi:10.1371/journal.pgen.1002768

23. Qin H, Cressy M, Li W, Coravos JS, Izzi SA, Dubnau J. Gamma neurons mediate dopaminergic input during aversive olfactory memory formation in Drosophila. Curr Biol. 2012;22: 608–614.

24. Shuai Y, Hirokawa A, Ai Y, Zhang M, Li W, Zhong Y. Dissecting neural pathways for forgetting in Drosophila olfactory aversive memory. Proc Natl Acad Sci U S A. 2015;112: E6663–72.

25. Yamagata N, Ichinose T, Aso Y, Plaçais P-Y, Friedrich AB, Sima RJ, et al. Distinct dopamine neurons mediate reward signals for short- and long-term memories. Proc Natl Acad Sci U S A. 2015;112: 578–583.

26. Aso Y, Rubin GM. Dopaminergic neurons write and update memories with cell-type-specific rules. Elife. 2016;5. doi:10.7554/eLife.16135

27. Berry JA, Phan A, Davis RL. Dopamine Neurons Mediate Learning and Forgetting through Bidirectional Modulation of a Memory Trace. Cell Rep. 2018;25: 651–662.e5.

28. Aso Y, Ray RP, Long X, Bushey D, Cichewicz K, Ngo T-T, et al. Nitric oxide acts as a cotransmitter in a subset of dopaminergic neurons to diversify memory dynamics. Elife. 2019;8. doi:10.7554/eLife.49257

29. Siju KP, Bräcker LB, Grunwald Kadow IC. Neural mechanisms of context-dependent processing of CO2 avoidance behavior in fruit flies. Fly . 2014;8: 68–74.

30. Dolan M-J, Belliart-Guérin G, Bates AS, Frechter S, Lampin-Saint-Amaux A, Aso Y, et al. Communication from Learned to Innate Olfactory Processing Centers Is Required for Memory Retrieval in Drosophila. Neuron. 2018;100: 651–668.e8.

31. Dolan M-J, Frechter S, Bates AS, Dan C, Huoviala P, Roberts RJ, et al. Neurogenetic dissection of the Drosophila lateral horn reveals major outputs, diverse behavioural functions, and interactions with the mushroom body. Elife. 2019;8. doi:10.7554/eLife.43079

32. Vogt K, Schnaitmann C, Dylla KV, Knapek S, Aso Y, Rubin GM, et al. Shared mushroom body circuits underlie visual and olfactory memories in Drosophila | eLife. Elife. 2014;3: e02395.

33. Ueno T, Tomita J, Tanimoto H, Endo K, Ito K, Kume S, et al. Identification of a dopamine pathway that regulates sleep and arousal in Drosophila. Nat Neurosci. 2012;15: 1516–1523.

34. Sitaraman D, Aso Y, Rubin GM, Nitabach MN. Control of Sleep by Dopaminergic Inputs to the Drosophila Mushroom Body. Front Neural Circuits. 2015;9: 73.

35. Bang S, Hyun S, Hong S-T, Kang J, Jeong K, Park J-J, et al. Dopamine signalling in mushroom bodies regulates temperature-preference behaviour in Drosophila. PLoS Genet. 2011;7: e1001346.

36. Azanchi R, Kaun KR, Heberlein U. Competing dopamine neurons drive oviposition choice for ethanol in Drosophila. Proc Natl Acad Sci U S A. 2013;110: 21153–21158.

37. Lim J, Fernandez AI, Hinojos SJ, Aranda GP, James J, Seong C-S, et al. The mushroom body D1 dopamine receptor controls innate courtship drive. Genes Brain Behav. 2018;17: 158–167.

38. Aimon S, Katsuki T, Jia T, Grosenick L, Broxton M, Deisseroth K, et al. Fast near-whole-brain imaging in adult Drosophila during responses to stimuli and behavior. PLoS Biol. 2019;17: e2006732.

39. Liu L, Wolf R, Ernst R, Heisenberg M. Context generalization in Drosophila visual learning requires the mushroom bodies. Nature. 1999;400: 753–756.

40. Zhang K, Guo JZ, Peng Y, Xi W, Guo A. Dopamine-mushroom body circuit regulates saliency-based decision-making in Drosophila. Science. 2007;316: 1901–1904.

41. van Swinderen B, McCartney A, Kauffman S, Flores K, Agrawal K, Wagner J, et al. Shared visual attention and memory systems in the Drosophila brain. PLoS One. 2009;4: e5989.

42. Manjila SB, Kuruvilla M, Ferveur J-F, Sane SP, Hasan G. Extended Flight Bouts Require Disinhibition from GABAergic Mushroom Body Neurons. Curr Biol. 2019;29: 283–293.e5.

43. Tsao C-H, Chen C-C, Lin C-H, Yang H-Y, Lin S. Drosophila mushroom bodies integrate hunger and satiety signals to control innate food-seeking behavior. Elife. 2018;7. doi:10.7554/eLife.35264

44. Sayin S, De Backer J-F, Siju KP, Wosniack ME, Lewis LP, Frisch L-M, et al. A Neural Circuit Arbitrates between Persistence and Withdrawal in Hungry Drosophila. Neuron. 2019;104: 544–558.e6.

45. Zolin A, Cohn R, Pang R, Siliciano AF, Fairhall AL, Ruta V. Context-dependent representations of movement in Drosophila dopaminergic reinforcement pathways. Nat Neurosci. 2021;24: 1555–1566.

46. Hattori D, Aso Y, Swartz KJ, Rubin GM, Abbott LF, Axel R. Representations of Novelty and Familiarity in a Mushroom Body Compartment. Cell. 2017;169: 956–969.e17.

47. Colomb J, Kaiser L, Chabaud M-A, Preat T. Parametric and genetic analysis of Drosophila appetitive long-term memory and sugar motivation. Genes Brain Behav. 2009;8: 407–415.

48. Landayan D, Feldman DS, Wolf FW. Satiation state-dependent dopaminergic control of foraging in Drosophila. Sci Rep. 2018;8: 5777.

49. Grunwald Kadow IC. State-dependent plasticity of innate behavior in fruit flies. Curr Opin Neurobiol. 2019;54: 60–65.

50. Aso Y, Hattori D, Yu Y, Johnston RM, Iyer NA, Ngo T-TB, et al. The neuronal architecture of the mushroom body provides a logic for associative learning. Elife. 2014;3: e04577.

51. Takemura S-Y, Aso Y, Hige T, Wong A, Lu Z, Shan Xu C, et al. A connectome of a learning and memory center in the adult Drosophila brain. Elife. 2017;6. doi:10.7554/elife.26975

52. Aso Y, Sitaraman D, Ichinose T, Kaun KR, Vogt K, Belliart-Guérin G, et al. Mushroom body output neurons encode valence and guide memory-based action selection in Drosophila. Elife. 2014;3: e04580.

53. Hige T, Aso Y, Modi MN, Rubin GM, Turner GC. Heterosynaptic Plasticity Underlies Aversive Olfactory Learning in Drosophila. Neuron. 2015;88: 985–998.

54. Cohn R, Morantte I, Ruta V. Coordinated and Compartmentalized Neuromodulation Shapes Sensory Processing in Drosophila. Cell. 2015;163: 1742–1755.

55. Owald D, Waddell S. Olfactory learning skews mushroom body output pathways to steer behavioral choice in Drosophila. Curr Opin Neurobiol. 2015;35: 178–184.

56. Jacob PF, Vargas-Gutierrez P, Okray Z, Vietti-Michelina S, Felsenberg J, Waddell S. Prior experience conditionally inhibits the expression of new learning in Drosophila. Curr Biol. 2021;31: 3490–3503.e3.

57. Siju KP, Štih V, Aimon S, Gjorgjieva J, Portugues R, Grunwald Kadow IC. Valence and State-Dependent Population Coding in Dopaminergic Neurons in the Fly Mushroom Body. Curr Biol. 2020;30: 2104–2115.e4.

58. Jovanoski KD, Duquenoy L, Mitchell J, Kapoor I, Treiber CD, Croset V, et al. Dopaminergic systems create reward seeking despite adverse consequences. Nature. 2023. doi:10.1038/s41586-023-06671-8

59. Cheng KY, Colbath RA, Frye MA. Olfactory and Neuromodulatory Signals Reverse Visual Object Avoidance to Approach in Drosophila. Curr Biol. 2019;29: 2058–2065.e2.

60. Huetteroth W, Perisse E, Lin S, Klappenbach M, Burke C, Waddell S. Sweet taste and nutrient value subdivide rewarding dopaminergic neurons in Drosophila. Curr Biol. 2015;25: 751–758.

61. Otto N, Pleijzier MW, Morgan IC, Edmondson-Stait AJ, Heinz KJ, Stark I, et al. Input Connectivity Reveals Additional Heterogeneity of Dopaminergic Reinforcement in Drosophila. Curr Biol. 2020;30: 3200–3211.e8.

62. Stern U, Srivastava H, Chen H-L, Mohammad F, Claridge-Chang A, Yang C-H. Learning a Spatial Task by Trial and Error in Drosophila. Curr Biol. 2019;29: 2517–2525.e5.

63. Mishra A, Cronley P, Ganesan M, Schulz DJ, Zars T. Dopaminergic neurons can influence heat-box place learning in Drosophila. J Neurogenet. 2020;34: 115–122.

64. Lesar A, Tahir J, Wolk J, Gershow M. Switch-like and persistent memory formation in individual Drosophila larvae. Elife. 2021;10. doi:10.7554/eLife.70317

65. Rohrsen C, Kumpf A, Semiz K, Aydin F, deBivort B, Brembs B. Pain is so close to pleasure: the same dopamine neurons can mediate approach and avoidance in Drosophila. bioRxiv. 2021. p. 2021.10.04.463010. doi:10.1101/2021.10.04.463010

66. Klapoetke NC, Murata Y, Kim SS, Pulver SR, Birdsey-Benson A, Cho YK, et al. Independent optical excitation of distinct neural populations. Nat Methods. 2014;11: 338–346.

67. Nässel DR, Elekes K. Aminergic neurons in the brain of blowflies and Drosophila: dopamine- and tyrosine hydroxylase-immunoreactive neurons and their relationship with putative histaminergic neurons. Cell Tissue Res. 1992;267: 147–167.

68. Pfeiffer BD, Jenett A, Hammonds AS, Ngo T-TB, Misra S, Murphy C, et al. Tools for neuroanatomy and neurogenetics in Drosophila. Proc Natl Acad Sci U S A. 2008;105: 9715–9720.

69. Jenett A, Rubin GM, Ngo T-TB, Shepherd D, Murphy C, Dionne H, et al. A GAL4-driver line resource for Drosophila neurobiology. Cell Rep. 2012;2: 991–1001.

70. Perisse E, Burke C, Huetteroth W, Waddell S. Shocking revelations and saccharin sweetness in the study of Drosophila olfactory memory. Curr Biol. 2013;23: R752–63.

71. Ho J, Tumkaya T, Aryal S, Choi H, Claridge-Chang A. Moving beyond P values: data analysis with estimation graphics. Nat Methods. 2019 [cited 20 Jun 2019]. doi:10.1038/s41592-019-0470-3

72. Dubnau J, Grady L, Kitamoto T, Tully T. Disruption of neurotransmission in Drosophila mushroom body blocks retrieval but not acquisition of memory. Nature. 2001;411: 476–480.

73. McGuire SE, Le PT, Davis RL. The role of Drosophila mushroom body signaling in olfactory memory. Science. 2001;293: 1330–1333.

74. Govorunova EG, Sineshchekov OA, Janz R, Liu X, Spudich JL. Natural light-gated anion channels: A family of microbial rhodopsins for advanced optogenetics. Science. 2015;349: 647–650.

75. Mohammad F, Stewart JC, Ott S, Chlebikova K, Chua JY, Koh T-W, et al. Optogenetic inhibition of behavior with anion channelrhodopsins. Nat Methods. 2017. doi:10.1038/nmeth.4148

76. Liu C, Plaçais P-Y, Yamagata N, Pfeiffer BD, Aso Y, Friedrich AB, et al. A subset of dopamine neurons signals reward for odour memory in Drosophila. Nature. 2012;488: 512–516.

77. Sun H, Nishioka T, Hiramatsu S, Kondo S, Amano M, Kaibuchi K, et al. Dopamine Receptor Dop1R2 Stabilizes Appetitive Olfactory Memory through the Raf/MAPK Pathway in Drosophila. J Neurosci. 2020;40: 2935–2942.

78. Dietzl G, Chen D, Schnorrer F, Su K-C, Barinova Y, Fellner M, et al. A genome-wide transgenic RNAi library for conditional gene inactivation in Drosophila. Nature. 2007;448: 151–156.

79. Molinoff PB, Axelrod J. Biochemistry of catecholamines. Annu Rev Biochem. 1971;40: 465–500.

80. Bainton RJ, Tsai LT, Singh CM, Moore MS, Neckameyer WS, Heberlein U. Dopamine modulates acute responses to cocaine, nicotine and ethanol in Drosophila. Curr Biol. 2000;10: 187–194.

81. Neckameyer WS. Multiple roles for dopamine in Drosophila development. Dev Biol. 1996;176: 209–219.

82. Montague SA, Baker BS. Memory Elicited by Courtship Conditioning Requires Mushroom Body Neuronal Subsets Similar to Those Utilized in Appetitive Memory. PLoS One. 2016;11: e0164516.

83. Dag U, Lei Z, Le JQ, Wong A, Bushey D, Keleman K. Neuronal reactivation during post-learning sleep consolidates long-term memory in Drosophila. Elife. 2019;8. doi:10.7554/eLife.42786

84. Handler A, Graham TGW, Cohn R, Morantte I, Siliciano AF, Zeng J, et al. Distinct Dopamine Receptor Pathways Underlie the Temporal Sensitivity of Associative Learning. Cell. 2019;178: 60–75.e19.

85. Jacob PF, Waddell S. Spaced Training Forms Complementary Long-Term Memories of Opposite Valence in Drosophila. Neuron. 2020. doi:10.1016/j.neuron.2020.03.013

86. Borenstein M, Hedges LV, Higgins JPT, Rothstein HR. Introduction to Meta-Analysis. Wiley; 2009.

87. Aguilar JI, Dunn M, Mingote S, Karam CS, Farino ZJ, Sonders MS, et al. Neuronal Depolarization Drives Increased Dopamine Synaptic Vesicle Loading via VGLUT. Neuron. 2017;95: 1074–1088.e7.

88. Owald D, Felsenberg J, Talbot CB, Das G, Perisse E, Huetteroth W, et al. Activity of defined mushroom body output neurons underlies learned olfactory behavior in Drosophila. Neuron. 2015;86: 417–427.

89. Cognigni P, Felsenberg J, Waddell S. Do the right thing: neural network mechanisms of memory formation, expression and update in Drosophila. Curr Opin Neurobiol. 2018;49: 51–58.

90. Boto T, Louis T, Jindachomthong K, Jalink K, Tomchik SM. Dopaminergic modulation of cAMP drives nonlinear plasticity across the Drosophila mushroom body lobes. Curr Biol. 2014;24: 822–831.

91. Takemura S-Y, Aso Y, Hige T, Wong A, Lu Z, Xu CS, et al. A connectome of a learning and memory center in the adult Drosophila brain. Elife. 2017;6. doi:10.7554/eLife.26975

92. Tanaka NK, Tanimoto H, Ito K. Neuronal assemblies of the Drosophila mushroom body. J Comp Neurol. 2008;508: 711–755.

93. Chiang A-S, Lin C-Y, Chuang C-C, Chang H-M, Hsieh C-H, Yeh C-W, et al. Three-dimensional reconstruction of brain-wide wiring networks in Drosophila at single-cell resolution. Curr Biol. 2011;21: 1–11.

94. Simões JM, Levy JI, Zaharieva EE, Vinson LT, Zhao P, Alpert MH, et al. Robustness and plasticity in Drosophila heat avoidance. Nat Commun. 2021;12: 2044.

95. Li F, Lindsey JW, Marin EC, Otto N, Dreher M, Dempsey G, et al. The connectome of the adult Drosophila mushroom body provides insights into function. Elife. 2020;9. doi:10.7554/eLife.62576

96. Yildizoglu T, Weislogel J-M, Mohammad F, Chan ES-Y, Assam PN, Claridge-Chang A. Estimating Information Processing in a Memory System: The Utility of Meta-analytic Methods for Genetics. PLoS Genet. 2015;11: e1005718.

97. Vogt K, Aso Y, Hige T, Knapek S, Ichinose T, Friedrich AB, et al. Direct neural pathways convey distinct visual information to Drosophila mushroom bodies. Elife. 2016;5. doi:10.7554/eLife.14009

98. Berry JA, Cervantes-Sandoval I, Chakraborty M, Davis RL. Sleep Facilitates Memory by Blocking Dopamine Neuron-Mediated Forgetting. Cell. 2015;161: 1656–1667.

99. Fuenzalida-Uribe N, Campusano JM. Unveiling the Dual Role of the Dopaminergic System on Locomotion and the Innate Value for an Aversive Olfactory Stimulus in Drosophila. Neuroscience. 2018;371: 433–444.

100. Berridge KC, Kringelbach ML. Pleasure systems in the brain. Neuron. 2015;86: 646–664.

101. Llinas RR. I of the Vortex: From Neurons to Self. MIT Press; 2001.

102. Croset V, Treiber CD, Waddell S. Cellular diversity in the Drosophila midbrain revealed by single-cell transcriptomics. Elife. 2018;7. doi:10.7554/eLife.34550

103. Yamazaki D, Maeyama Y, Tabata T. Combinatory Actions of Co-transmitters in Dopaminergic Systems Modulate Drosophila Olfactory Memories. J Neurosci. 2023;43: 8294–8305.

104. Sinakevitch I, Grau Y, Strausfeld NJ, Birman S. Dynamics of glutamatergic signaling in the mushroom body of young adult Drosophila. Neural Dev. 2010;5: 10.

105. Miyashita T, Murakami K, Kikuchi E, Ofusa K, Mikami K, Endo K, et al. Glia transmit negative valence information during aversive learning in *Drosophila*. Science. 2023;382: eadf7429.

106. Sabandal JM, Sabandal PR, Kim Y-C, Han K-A. Concerted Actions of Octopamine and Dopamine Receptors Drive Olfactory Learning. J Neurosci. 2020;40: 4240–4250.

107. Iliadi KG, Iliadi N, Boulianne GL. Drosophila mutants lacking octopamine exhibit impairment in aversive olfactory associative learning. Eur J Neurosci. 2017;46: 2080–2087.

108. Chuhma N, Zhang H, Masson J, Zhuang X, Sulzer D, Hen R, et al. Dopamine neurons mediate a fast excitatory signal via their glutamatergic synapses. J Neurosci. 2004;24: 972–981.

109. Stuber GD, Hnasko TS, Britt JP, Edwards RH, Bonci A. Dopaminergic terminals in the nucleus accumbens but not the dorsal striatum corelease glutamate. J Neurosci. 2010;30: 8229–8233.

110. Chuhma N, Mingote S, Moore H, Rayport S. Dopamine neurons control striatal cholinergic neurons via regionally heterogeneous dopamine and glutamate signaling. Neuron. 2014;81: 901–912.

111. Sherer LM, Catudio Garrett E, Morgan HR, Brewer ED, Sirrs LA, Shearin HK, et al. Octopamine neuron dependent aggression requires dVGLUT from dual-transmitting neurons. PLoS Genet. 2020;16: e1008609.

112. Yang C-H, Shih M-FM, Chang C-C, Chiang M-H, Shih H-W, Tsai Y-L, et al. Additive Expression of Consolidated Memory through Drosophila Mushroom Body Subsets. PLoS Genet. 2016;12: e1006061.

113. Perkins LA, Holderbaum L, Tao R, Hu Y, Sopko R, McCall K, et al. The Transgenic RNAi Project at Harvard Medical School: Resources and Validation. Genetics. 2015;201: 843–852.

114. Witten IB, Steinberg EE, Lee SY, Davidson TJ, Zalocusky KA, Brodsky M, et al. Recombinase-driver rat lines: tools, techniques, and optogenetic application to dopamine-mediated reinforcement. Neuron. 2011;72: 721–733.

115. Danjo T, Yoshimi K, Funabiki K, Yawata S, Nakanishi S. Aversive behavior induced by optogenetic inactivation of ventral tegmental area dopamine neurons is mediated by dopamine D2 receptors in the nucleus accumbens. Proc Natl Acad Sci U S A. 2014;111: 6455–6460.

116. Schultz W, Dayan P, Montague PR. A neural substrate of prediction and reward. Science. 1997;275: 1593–1599.

117. Kato S, Kaplan HS, Schrödel T, Skora S, Lindsay TH, Yemini E, et al. Global brain dynamics embed the motor command sequence of Caenorhabditis elegans. Cell. 2015;163: 656–669.

118. Ichinose T, Aso Y, Yamagata N, Abe A, Rubin GM, Tanimoto H. Reward signal in a recurrent circuit drives appetitive long-term memory formation. Elife. 2015;4: e10719.

119. Zhao X, Lenek D, Dag U, Dickson BJ, Keleman K. Persistent activity in a recurrent circuit underlies courtship memory in Drosophila. Elife. 2018;7. doi:10.7554/eLife.31425

120. Pavlowsky A, Schor J, Plaçais P-Y, Preat T. A GABAergic Feedback Shapes Dopaminergic Input on the Drosophila Mushroom Body to Promote Appetitive Long-Term Memory. Curr Biol. 2018;28: 1783–1793.e4.

121. Cumming G, Calin-Jageman R. Introduction to the New Statistics: Estimation, Open Science, and Beyond. Routledge; 2016.

122. Dulcis D, Lippi G, Stark CJ, Do LH, Berg DK, Spitzer NC. Neurotransmitter Switching Regulated by miRNAs Controls Changes in Social Preference. Neuron. 2017;95: 1319–1333.e5.

123. Dulcis D, Jamshidi P, Leutgeb S, Spitzer NC. Neurotransmitter switching in the adult brain regulates behavior. Science. 2013;340: 449–453.

124. Meng D, Li H-Q, Deisseroth K, Leutgeb S, Spitzer NC. Neuronal activity regulates neurotransmitter switching in the adult brain following light-induced stress. Proc Natl Acad Sci U S A. 2018;115: 5064–5071.

125. Buck SA, Steinkellner T, Aslanoglou D, Villeneuve M, Bhatte SH, Childers VC, et al. VGLUT modulates sex differences in dopamine neuron vulnerability to age-related neurodegeneration. bioRxiv. 2020. p. 2020.11.11.379008. doi:10.1101/2020.11.11.379008

126. Temasek Life Sciences Laboratories. Recipe for Drosophila media. In: Zenodo [Internet]. 28 Feb 2018. doi:10.5281/zenodo.1185451

127. Jung Y, Kennedy A, Chiu H, Mohammad F, Claridge-Chang A, Anderson DJ. Neurons that Function within an Integrator to Promote a Persistent Behavioral State in Drosophila. Neuron. 2020;105: 322–333.e5.

128. Zars T, Fischer M, Schulz R, Heisenberg M. Localization of a short-term memory in Drosophila. Science. 2000;288: 672–675.

129. Pitman JL, Huetteroth W, Burke CJ, Krashes MJ, Lai S-L, Lee T, et al. A pair of inhibitory neurons are required to sustain labile memory in the Drosophila mushroom body. Curr Biol. 2011;21: 855–861.

130. Kan L, Ott S, Joseph B, Park ES, Dai W, Kleiner RE, et al. A neural m6A/Ythdf pathway is required for learning and memory in Drosophila. Nat Commun. 2021;12: 1458.

131. Tully T, Quinn WG. Classical conditioning and retention in normal and mutant Drosophila melanogaster. J Comp Physiol A. 1985;157: 263–277.

132. DiCiccio TJ, Efron B. Bootstrap Confidence Intervals. Stat Sci. 1996;11: 189–228.

133. Tumkaya T, Ott S, Claridge-Chang A. Systematic review of Drosophila short-term-memory genetics: Meta-analysis reveals robust reproducibility. Neurosci Biobehav Rev. 2018. doi:10.1016/j.neubiorev.2018.07.016

134. Altman D, Machin D, Bryant T, Gardner S. Statistics with confidence: confidence interval and statistical guidelines. Bristol: BMJ Books. 2000.

135. Efron B, Tibshirani RJ. An Introduction to the Bootstrap. CRC Press; 1994.

