## Supplementary File for "Dopamine neurons that inform *Drosophila* olfactory memory have distinct, acute functions driving attraction and aversion"

**Supplementary Figures**

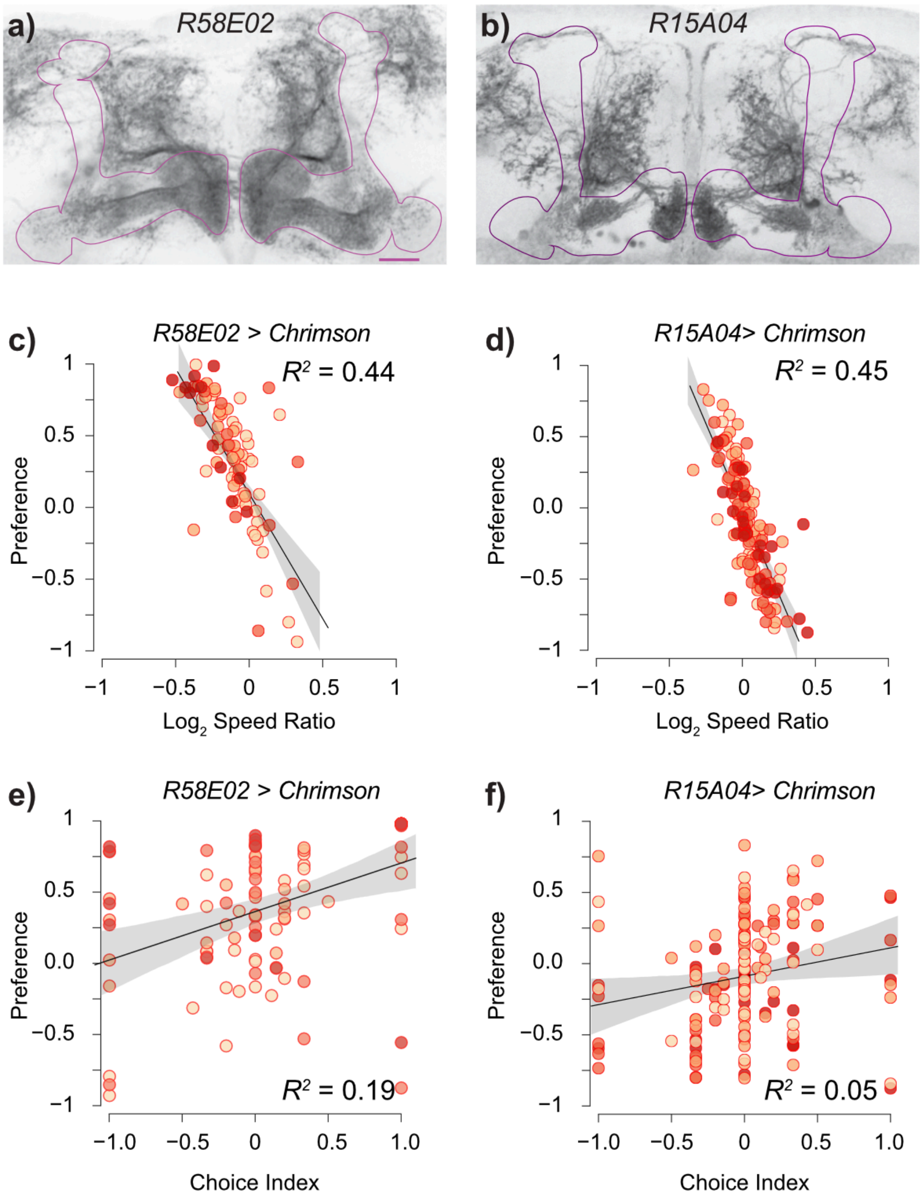


**Figure S1. Dopaminergic-line optogenetic preference is related to changes in walking speed.**
**a-b.** Maximum-intensity projections of genetically-stained mushroom bodies show innervation by *R58E02* and *R15A04* in various MB neuropil zones. Dark pixels indicate α-GFP signal from the GFP tag on ACR1. *R58E02* has stained fibers across nearly all the horizontal lobes. *R15A04* sends fibers to four zones, α1, β2, β'1, and γ5. The MB lobes (purple line) were outlined using DLG-1 counterstain, not shown here (see Methods for details)—scale bar: 20 µm. See also Movies S1 and S3.
**c.** Scatter plots and linear regression of *R58E02>Chr* preference indices (PIs) and their light/dark $\log_2$ speed ratios. Each point indicates metrics for a single fly. The two metrics are negatively correlated: $R^2_{adj}$ = 0.44 (95 CI, 0.24, 0.62, $P = 2.9 \times 10^{-12}$). $N$ = 86. Each point indicates the $\log_2$ speed ratio and PI of a single fly. Flies were assayed at 1.3, 5, 22, and 70 µW/mm$^2$.
**d.** There is an inverse relationship between *R15A04>Chr* PIs and speed ratios; $R^2_{adj}$ = 0.45 (95 CI, 0.3, 0.6, $P = 1.2 \times 10^{-20}$).
**e-f.** No correlation between PI and choice index was observed in *R58E02>Chr* ($R^2_{adj}$ = 0.01 (95 CI, 0.00, 0.07, $P = 3.4 \times 10^{-1}$) *and R515A04>Chr* ($R^2_{adj}$ = 0.04 (95 CI, 0.00, 0.05, $P = 4.1 \times 10^{-1}$)flies; . Each point indicates metrics for a single fly. Flies were screened at 1.3, 5, 22, and 70 µW/mm$^2$.

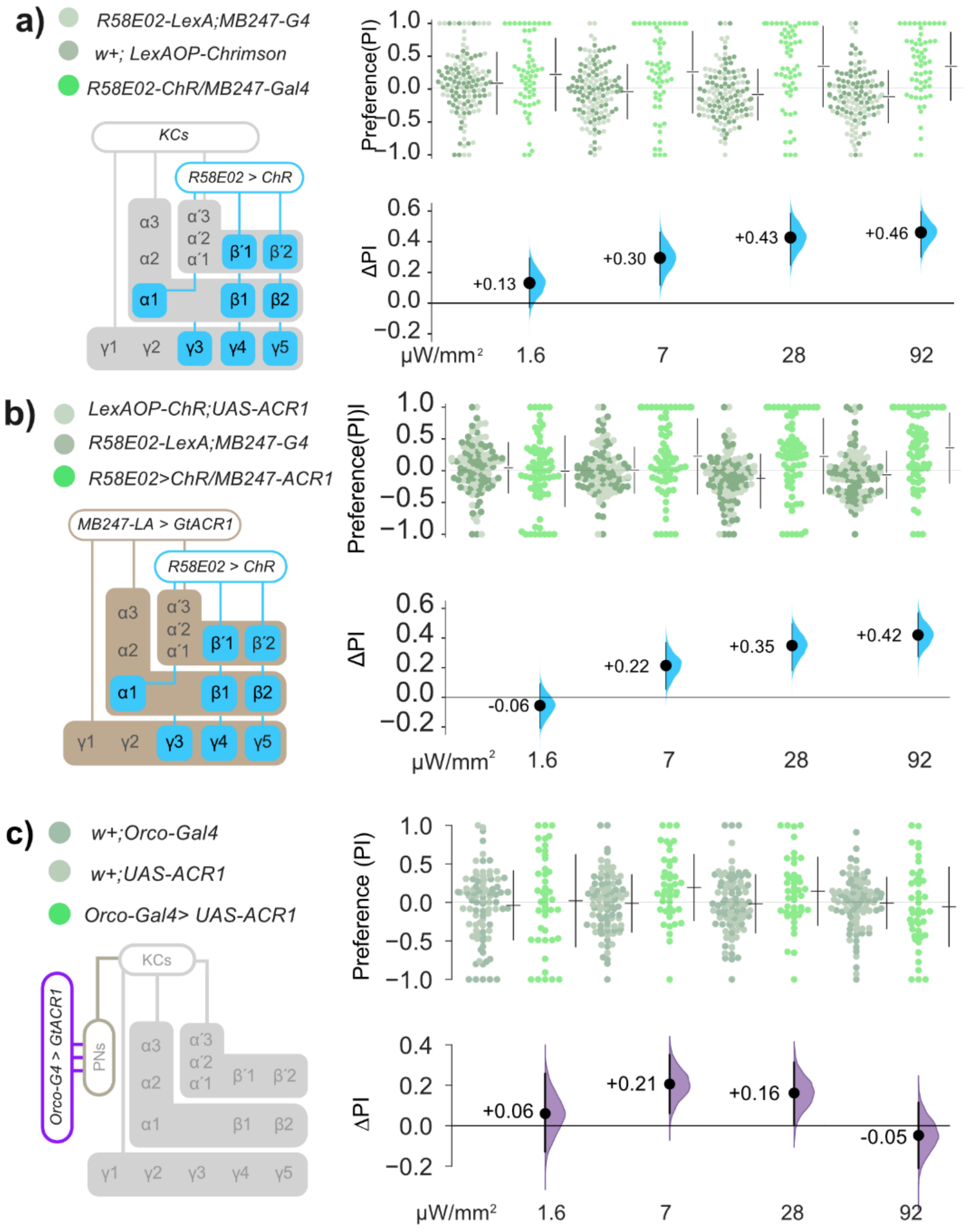


**Figure S2. Optogenetic PAM DAN valence does not require olfactory-system function.**

**a.** *R58E02-LexA>LexAop-Chr* flies are attracted to green light. The previous green-light optogenetic activation experiment (**Fig. 3a**) was replicated using the *R58E02-LexA* driver (instead of *R58E02-Gal4*) and reproduced the attraction phenotype.

**b.** In *R58E02>Chr/MB247>ACR1* flies, attraction to DAN self-activation is largely unaffected by simultaneous KC opto-inhibition (**Fig. 3b**). The green-light dual optogenetic experiment was replicated with *R58E02-LexA* and *MB247-Gal4* drivers; it reproduced the outcome observed in the previous experiment: valence is unaffected by inhibition of the *MB247* cells.

**c.** *Orco>ACR1* flies display moderate attraction to green light at two intermediate intensities (7 and 28 μW/mm²), but not at 92 μW/mm².

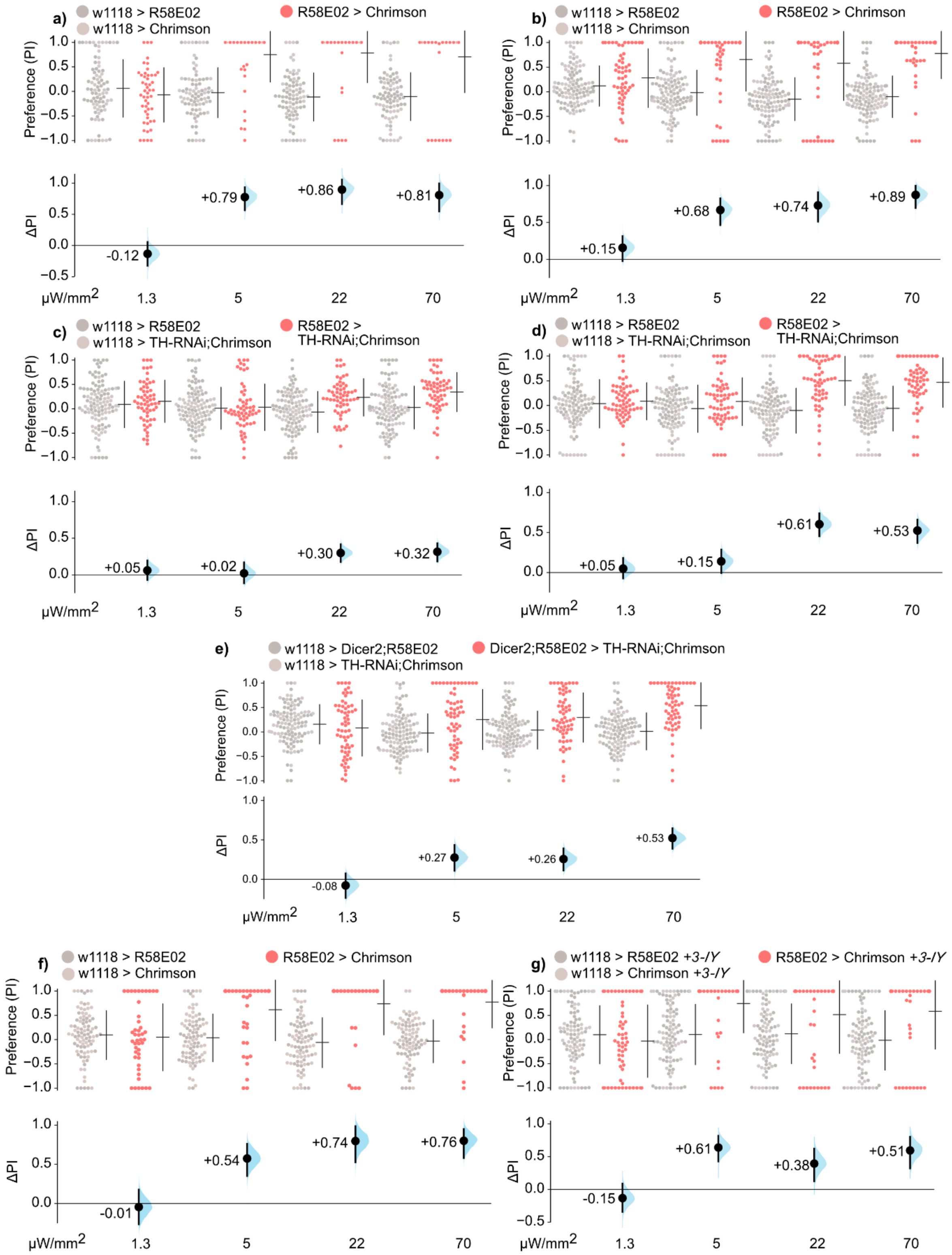


**Figure S3. Drug-induced and enhanced RNAi-mediated depletion of dopamine results in partially reduced valence.**

**a-b**. Additional replicates of *R58E02>Chrimson* used in **Fig. 4i** in the main text (see Methods for details). Data from **Fig. S3e** is also used in column 1 of main text **Fig. 5a**.

**c-d.** Additional replicates of *R58E02>TH-RNAi;Chrimson* used in main text **Fig. 4i** in the main text (see Methods for details).

**e.** Enhancement of RNAi-mediated TH knockdown via the overexpression of *Dicer2* resulted in valence at 22 μW/mm$^2$ of ΔPI = +0.26 [95CI +0.11, +0.39]; valence at 70 μW/mm$^2$ was ΔPI = +0.53 [95CI +0.39, +0.66].

**f-g.** Depletion of dopamine *via* feeding with 3-iodotyrosine (10 mg/mL 3-IY) resulted in a moderate reduction of valence. At 70 μW/mm$^2$, *R58E02>Chr* flies that were not fed 3-IY displayed valence of ΔPI = +0.76 [95CI +0.54, +0.92], whereas flies that were fed 3-IY displayed valence of ΔPI = +0.51 [95CI +0.25, +0.72]. The data from **Fig. S3f–g** is also used in columns 1–2 of the main text **Fig. 5b.**

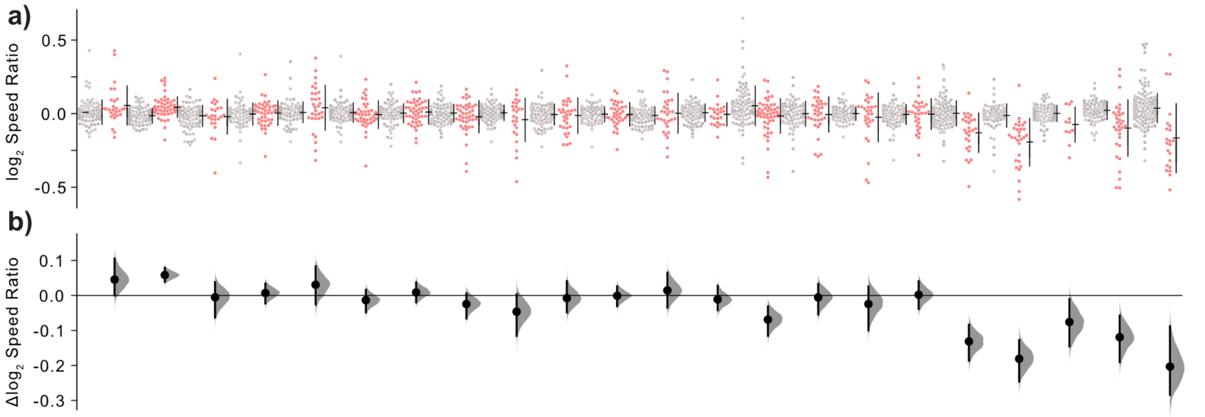


| Gal4 | R15A04 | NP5272 | 441B | 040B | 025B | 194B | 188B | 315C | 196B | 043B | 063B | 045B | 316B | 032B | 312B | 042B | 109B | 047B | 299B | 56B | 213B | R58E02 |
|---|---|---|---|---|---|---|---|---|---|---|---|---|---|---|---|---|---|---|---|---|---|---|
| PAM Type | M | 03 | 12 | M | 13,14 | M | 13,M | 01 | M | 11,M | 10 | 10,11 | M | 06,M | 07,08 | M | 01,02 | 05,M | 11,M | 05,06 | 10,M | M |

**Figure S4. Corresponding $\log_2$ speed ratios for PAM-DAN valence screen.**

**a-b.** Δ $\text{Log}_2$ speed ratios for the 22 PAM-DAN lines in our valence screen (**Fig. 6a-b**). Overall, lines with negative valence had positive speed ratios, while lines with positive valence had negative speed ratios. See **Table S1** for effect sizes. The table shows the PAM cell types where each driver expresses most strongly, and M denotes multiple cell types (DOI: 10.5281/zenodo.7239106). See **Table S2** for further details on driver expression.

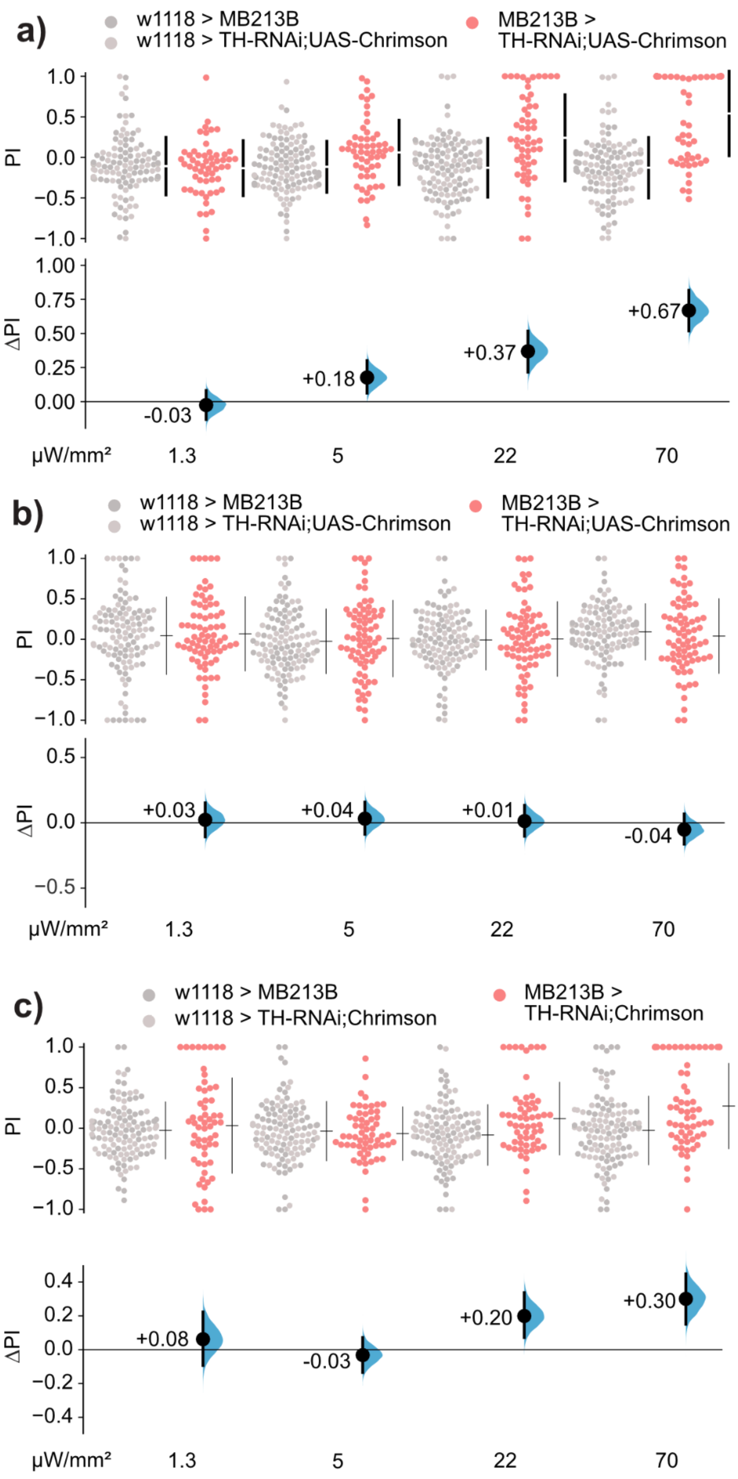


**Figure S5. TH knockdown in PAM-β cells led to variable reductions in effect size.**
**a-c.** Replicates of MB213B > TH-RNAi; Chrimson experiments showed varying effect sizes, necessitating meta-analysis. These panels correspond to Replicates 1, 2 and 3, respectively, in **Fig. 6e-f**.

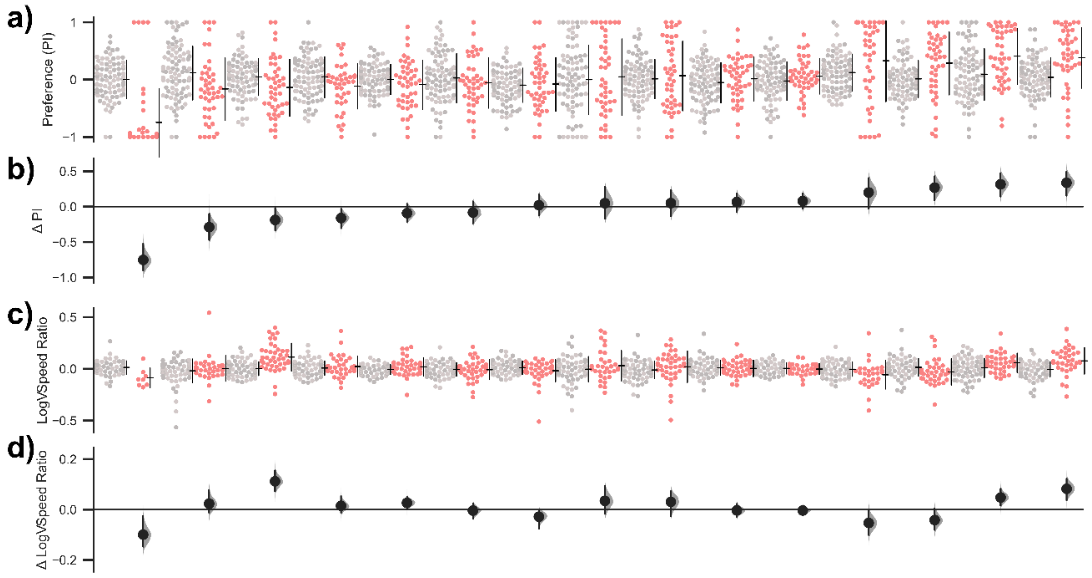


| MBON-Gal4 | VT999036 | 434B | 77B | R53C03 | R76B09 | 298B | 93C | R52G04 | 011B | 57B | VT41043 | VT49126 | 399C | 112C | 083C |
|---|---|---|---|---|---|---|---|---|---|---|---|---|---|---|---|
| Cell types | 20, 21 | 05, 06 | 12 | 05 | 05 | 05 | 13, 14 | M | M | 10 | 03 | 11 | 02 | 11 | 08, 09 |

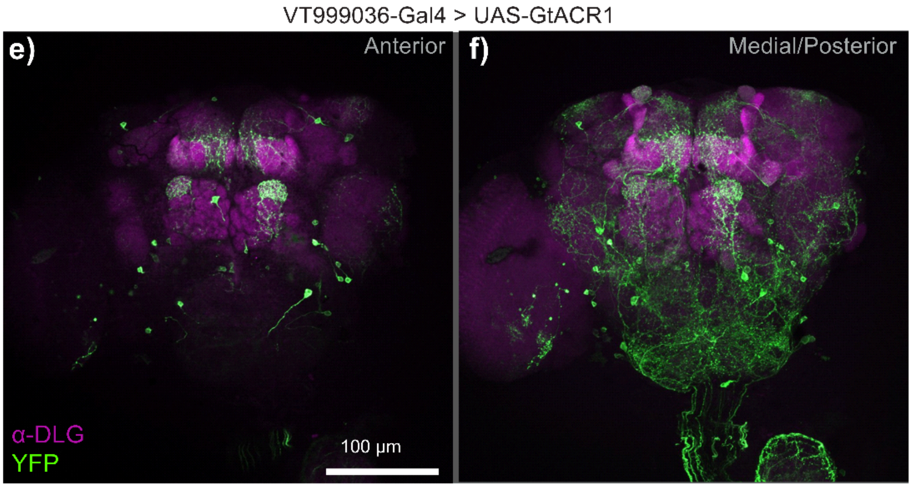


**Figure S6. In MBONs output synaptic zones, dopamine signaling is dispensable.**

**a-b.** An optogenetic valence screen for affective output zones using 15 MBONs related to split-Gal4 and Gal4 drivers. The data in panel **b** is a duplicate of **Fig. 7a**.

**c-d.** Δ $\text{Log}_2$ speed ratios for the 15 MBON lines in the optogenetic valence screen. See **Table S1** for effect sizes. The matrix key below indicates the drivers used and the corresponding MBON cell types in which they express; see **Table S2** for details.

**e-f.** Immunostaining of YFP in MBONs labeled by VT999036-Gal4. See also **Movie S6**.

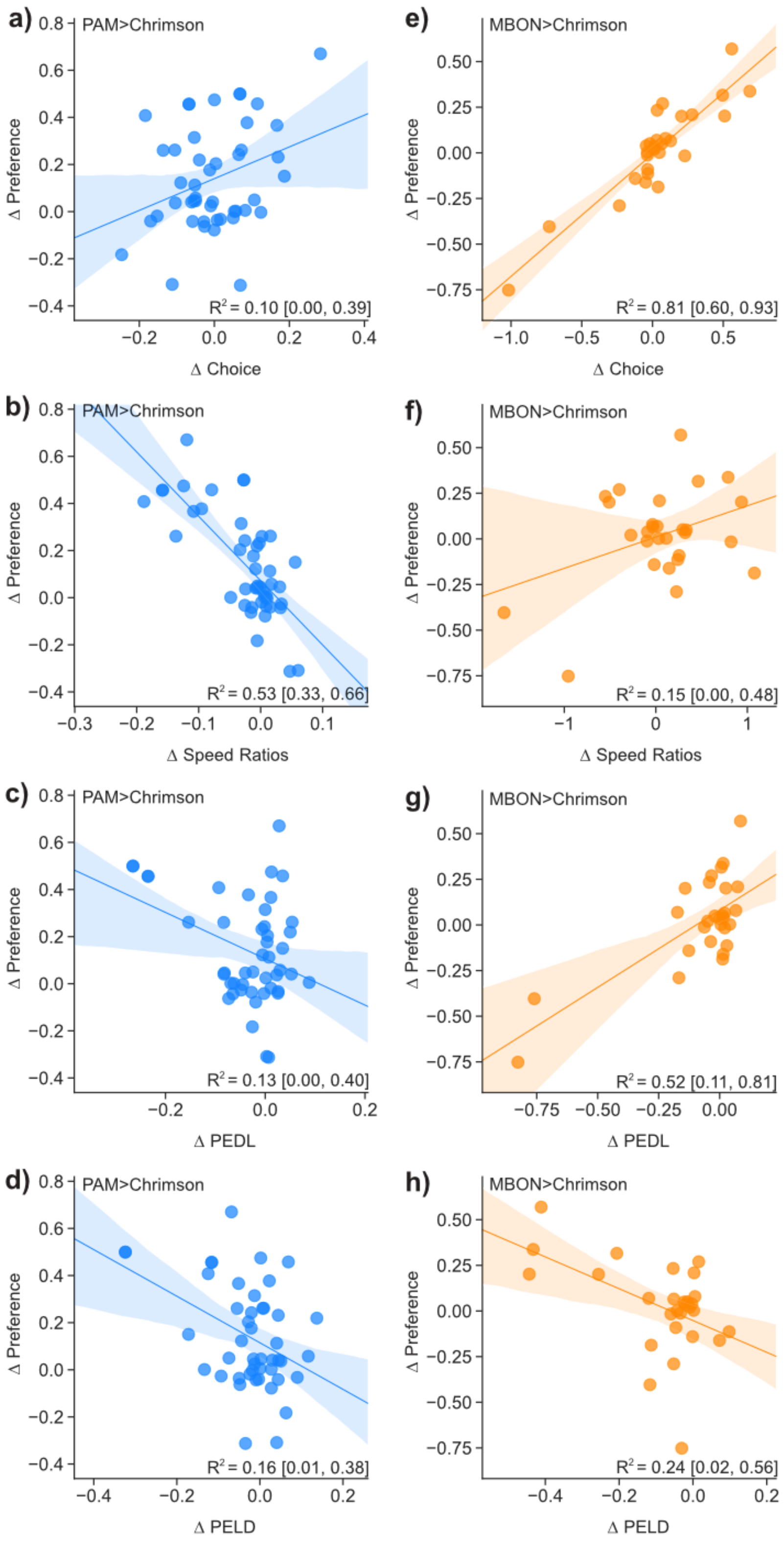


**Figure S7. Regression analyses of DAN and MBON drivers' behavior metrics.**

**a-d**. Scatter plots relating valence with Δchoice (ΔChI), Δspeed ratio, Δproportion of dark→light exits (ΔPEDL) and Δproportion of dark→light exits (ΔPELD) in the DAN lines.

**e-h.** Scatter plots relating valence to Δchoice, Δspeed ratio, Δproportion of dark→light exits (ΔPEDL) and Δproportion of dark→light exits (ΔPELD) for the MBON lines from Fig. 7a and S7a. The data are obtained from the two bright illumination intensities (22 and 70 μW/mm$^2$).

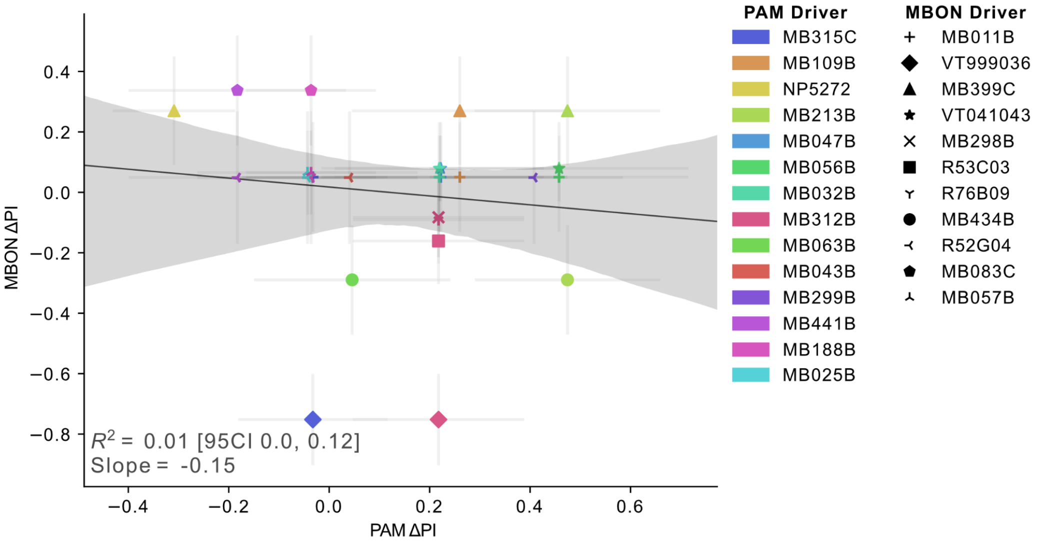


**Figure S8. No correlation between valence for co-zonal PAM and MBON activation.**
Valence scores from the PAM and MBON screens were compared for driver lines with similar projection patterns in the mushroom-body lobes. Each dot indicates the two valence scores from a pair of PAM:MBON drivers with major staining in identical neuropil zones. Error bars are the confidence intervals of the valence (ΔPI) scores. For each pair, the PAM driver is denoted by color, while the MBON driver is denoted by marker style.

### Supplementary Tables

**Table S1. Detailed genotypes and effect sizes**

Genotypes and effect sizes of experiments in each figure. Semicolons indicate different chromosomes, commas indicate different transgenes. Compound genotypes are stated for split-Gal4 drivers, with the common line designation (e.g. MB213B) in parentheses.

**Table S2. PAM and MBON types**

A catalog of published information about the drivers used in the PAM (Fig. 5a-b) and MBON (Fig. 4a, S4a) screens, the cell types they express in, and the lobes of the mushroom body in which each cell type's terminals can be found.

### Supplementary Movies

**Movie S1. Z-stack for *R58E02>ACR1* immunostaining.**

Full confocal stack of the PAM cluster in a representative *R58E02>ACR1* fly brain, corresponding to **Fig. S1a**. Green: YFP; Magenta: α-Dlg1; taken using 20× objective lens.

**Movie S2. *R58E02>Chr* flies walk slower in light**

Video of a 91-s-duration OSAR experiment, showing the dark phase and the first light epoch. *R58E02>Chr* flies slow or stop upon entering the light. The video is sped up to 1.33× realtime. Optogenetic light is turned on at t = 22 s and off at t= 106 s (video time). The light-border pattern was superimposed on the video (see Methods for details).

**Movie S3. Z-stack for *R15E02>ACR1* immunostaining.**

Full confocal stack of a subset of PAM neurons in a representative *R15E02>ACR1* fly brain, corresponding to **Fig. S1b**. Green: YFP; Red: α-Dlg1; taken using 20× objective lens.

**Movie S4. Z-stack for *R58E02>Chr* immunostaining.**

Full confocal stack of the PAM cluster in a representative *R58E02>Chr* fly brain, corresponding to **Fig. 3e**. Green: α-YFP; Red: α-TH; taken using 63× objective lens. There is high colocalization between YFP and TH signals.

**Movie S5. Z-stack for *R58E02>TH-RNAi;Chr* immunostaining.**

Full confocal stack of the PAM cluster in a representative *R58E02>TH-RNAi;Chr* fly brain, corresponding to **Fig. 3f**. Green: α-YFP; Red: α-TH; taken using 63× objective lens. The colocalization between YFP and TH signals is reduced.

**Movie S6. Z-stack for *VT999036>ACR1* immunostaining.**

Full confocal stack of a subset of MBON neurons in a representative *VT999036>ACR1* fly brain, corresponding to **Fig. S1b**. Green: YFP; Magenta: α-DLG; taken using 20× objective lens.

**Movie S7. *VT999036>Chr* flies avoid light**

*VT999036>Chr* flies avoid light using reversal and turning maneuvers. Timing is identical to **Movie S2**.

**Movie S8. Z-stack for *MB213B>Chr* immunostaining**

Full confocal stack of a representative *MB213B>Chr* fly brain, corresponding to **Fig. 5c**. Green: YFP; Magenta: α-DLG; taken using 20× objective lens.
